# Image3C: a multimodal image-based and label independent integrative method for single-cell analysis

**DOI:** 10.1101/603035

**Authors:** Alice Accorsi, Andrew C. Box, Robert Peuß, Christopher Wood, Alejandro Sánchez Alvarado, Nicolas Rohner

## Abstract

Image-based cell classification has become a common tool to identify phenotypic changes in cell populations. However, this methodology is limited to organisms possessing well characterized species-specific reagents (e.g., antibodies) that allow cell identification, clustering and convolutional neural network (CNN) training. In the absence of such reagents, the power of image-based classification has remained mostly off-limits to many research organisms. We have developed an image-based classification methodology we named Image3C (Image-Cytometry Cell Classification) that does not require species-specific reagents nor pre-existing knowledge about the sample. Image3C combines image-based flow cytometry with an unbiased, high-throughput cell cluster pipeline and CNN integration. Image3C exploits intrinsic cellular features and non-species-specific dyes to perform *de novo* cell composition analysis and to detect changes in cellular composition between different conditions. Therefore, Image3C expands the use of imaged-based analyses of cell population composition to research organisms in which detailed cellular phenotypes are unknown or for which species-specific reagents are not available.

**Impact statement:** Image3C analyzes cell populations through image-based clustering and neural network training, which allows single-cell analysis in research organisms devoid of species-specific reagents or pre-existing knowledge on cell phenotypes.

## Introduction

Single-cell analysis have proven crucial to our understanding of fundamental biological processes such as development, homeostasis, regeneration, aging and disease (Goolam et al., 2016; Kimmel et al., 2019; Pepe-Mooney et al., 2019; Philippeos et al., 2018; Tirosh et al., 2016). High-throughput analyses of these and other biological processes at single cell-resolution require technologies capable of describing individual cells and subsequently clustering them based on similarities of features like morphology, cell surface protein expression or transcriptome profile. Recent advances in image-based cell profiling and single-cell RNA sequencing (scRNA-seq) allow quantification of differences between cell populations and comparisons of cell type composition between samples (Caicedo et al., 2017). Single-cell studies that use traditional research organisms (*e.g.,* mouse, rat or fruit fly) benefit from the availability of genomic platforms and established antibody libraries. However, the same cannot be said for a growing number of important, yet understudied research organism lacking such reagents and whose biological interrogation would benefit immensely from single-cell analyses. In these cases, classical histochemical methods are often used to identify and characterize specific cells. Yet, the successful identification and enumeration of biologically meaningful cell types in such studies can be harmed by both the limited number and variety of cellular attributes (few features or low dynamic range) available for determination of cell types, and by observer bias when using traditional, hand-counting approaches (*e.g.*, hemocytometer and Giemsa stain) (van der Meer, Scott, & de Keijzer, 2004). These shortcomings, together with the lack of extensive knowledge on cell-specific phenotypes available for training or for *a priori* assumptions usually results in the underestimation of the complexity of cellular composition or interactions among cell types within tissues.

Automated classification of cells using convolutional neural networks (CNN, machine learning method specialized in image recognition and classification) has become a promising approach for accurate high-throughput cell analysis that is free from observer bias (Blasi et al., 2016; Eulenberg et al., 2017; Kobayashi et al., 2017; Lei et al., 2018; Nassar et al., 2019). To date, CNN-based automated clustering and classification techniques require pre-existing knowledge about the organism or cell type of interest *(e.g.,* cell specific morphological traits within an image set) or the availability of cell-specific reagents (*e.g.*, antibodies), or genomic sequence (*e.g.*, single-cell sequencing) (Baron et al., 2019; Blasi et al., 2016; Eulenberg et al., 2017; Kobayashi et al., 2017; Lei et al., 2018; Nassar et al., 2019). This means that to make effective use of artificial intelligence (AI) approaches for single-cell analysis, one must have information available to train the algorithm or for machine learning (ML) models, which often arises in the form of information gleaned from the use of reagents like antibodies. Research areas that rely on inter-species comparisons or studies on emerging research organisms would benefit from single cell-based analyses that do not require pre-existing knowledge of cell types *(i.e.,* which is required for training a CNN for example) and/or availability of antibodies or molecular databases. For example, within the interdisciplinary field of eco-immunology, a growing number of researchers is investigating immune system adaptation to different environments by studying immune cell compositions in diverse animals (Maizels & Nussey, 2013). Given the influences of immune cell composition on the immune system response of an organism (Kaczorowski et al., 2017), applying modern single cell analysis in eco-immunological research would substantially increase our knowledge about the plasticity and conservation of immune responses in a variety of different animals and conditions(Peuß et al., 2020).

To make sophisticated cellular composition analysis available to any research organism without the need for either pre-existing knowledge about the cell populations or species-specific reagents, we developed Image-Cytometry Cell Classification (Image3C). This method analyzes, visualizes, and quantifies, in a high-throughput and unbiased way, the composition of cell populations by using cell morphological traits and non-species-specific fluorescent probes (*e.g.*, nuclear staining or dyes for metabolic states that function well in a variety of organisms). By taking advantage of cell morphology and/or fluorescent dyes related to function or metabolic state, Image3C can analyze single cell suspensions derived from any experimental design, identify different cell types and cluster them. Once the cell types are identified by *de novo* clustering on cell intrinsic features, Image3C employs a CNN that uses these clusters as training sets, avoiding in this way user bias or manual classification. This produces a CNN-based cell classifier ‘machine’ used to quantify subsequently acquired image-based flow cytometry data and to compare cellular composition of samples across multiple experiments, in a high-throughput and unsupervised manner without the need for repeating time-consuming steps for *de novo* clustering. In comparison to existing label-free cell clustering methods, Image3C does not require initial antibody staining (Lippeveld et al., 2020; Nassar et al., 2019), pre-existing knowledge of specific cell morphology (Yakimov et al., 2019) and is not limited to a specific cellular phenotype (Blasi et al., 2016) for *a priori* identification of certain cell types. This makes Image3C extremely versatile and applicable to virtually any research organism and tissue from which dissociated single cells can be obtained. In sum, Image3C combines modern high-throughput data acquisition by image-based flow cytometry, advanced and unbiased clustering analysis, statistics to compare cellular compositions across different samples and a CNN classifier component to easily determine changes in cell composition across multiple experiments.

## Results and Discussion

### Image-Cytometry Cell Classification (Image3C)

Image3C is an imaging tool developed to study tissue composition at single-cell resolution in research organisms for which antibodies and pre-existing knowledge about cell types are not readily available. Image3C allows for high-throughput and unbiased analysis in scenarios where manual counting and observer-based cell identification are currently the only options. Image3C includes all the components required for compensating captured images, quantifying multiple features for each event, clustering the events, visualizing and exploring the data, training and using the CNN for analyzing subsequent samples and integrating multiple experiments (Figure 1 and S1).

**Figure 1:**
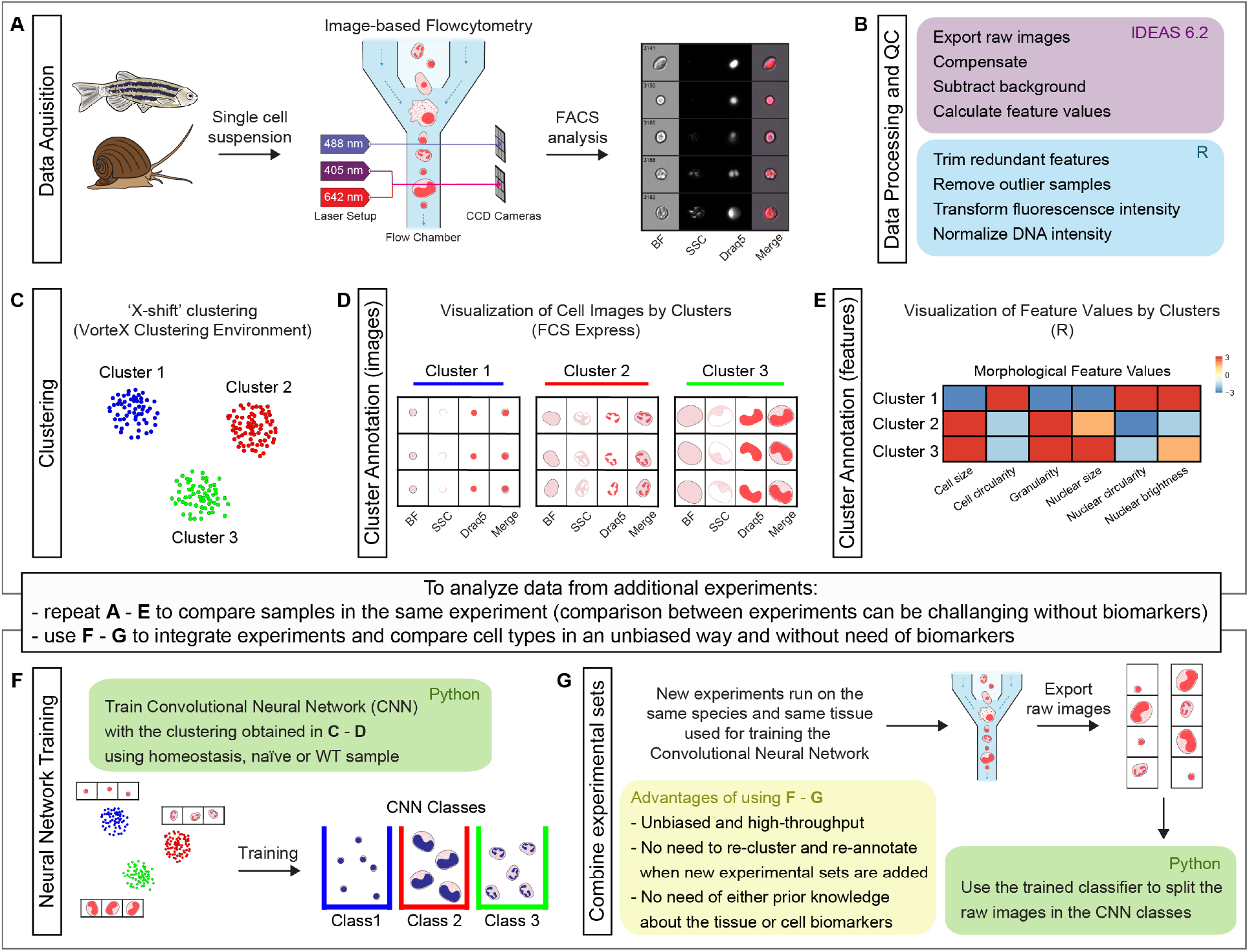
Schematic representation of Image3C workflow, a method for cell clustering based on morphological features. (A) A single cell suspension is prepared for image-based flow cytometric analyses. The cells can be labelled with any reagent working for the species of interest. The signal can highlight specific cell components *(e.g.,* nuclei), metabolic cell states or specific cell functions. The samples are run on the ImageStream®X Mark II and 10,000 nucleated and focused events are saved for each sample as individual raw images. (B) IDEAS software (Amnis Millipore) is used to open the raw images, compensate for correcting fluorescent spillover, subtract background and quantify values for intrinsic morphological and fluorescent features. R (or R studio) is used to calculate the correlation between features to allow to trim the features that are redundant with others. Samples that are outliers among replicates are also removed prior the final normalization of the fluorescence intensities. (C) Images are clustered based on morphological and fluorescent feature values and visualized as a Force Directed Layout (FDL) graph where each dot represents one event. (D) R integration in FCS Express software allows the visualization of cell images by clusters or specifically selected with a gate. This step allows to evaluate the morphological homogeneity of the clusters, determine if the number of clusters is appropriate and explore the phenotype/function of the cells based on visualization of individual channels. (E) Spearman’s correlation plot of feature values by clusters is one of the options available in Image3C for plotting integrated data. This heatmap shows feature similarities and differences between cells belonging to different clusters. (F) If new experiments are run and new data needs to be analyzed two approaches can be taken. 1. If the goal is comparing samples belonging to the same experiment (*e.g.*, treatment vs its control) the steps described so far from (A) to (E) can be re-applied to the new dataset including a statistical analysis to compare cluster relative abundance. This approach will produce a new set of clusters that will need to be re-annotated. Compare sets of clusters coming from multiple experiments and multiple rounds of analysis can be challenging without pre-existing knowledge of cell-type, clearly different morphologies or biomarkers that would allow to establish a unique correlation between clusters coming from different FDL graphs. 2. If the goal is integrating experiments and comparing cell type abundance between them, the use of steps (F) and (G) is suggested. A CNN classifier is trained using the images obtained from homeostasis, naïve or WT cells and already organized in clusters in an unbiased way through the first part of our method. This will generate a trained classifier with CNN classes based on FDL clusters. (G) This classifier is then used for deconvoluting data from new experimental sets and assigning each event to a CNN class with a given probability. This provides high-throughput, unsupervised and unbiased way to compare different experiment sets without the requirement for pre-existing knowledge about the tissue cell types, cell biomarkers or the need to cross-annotate cluster increasing the probability to introduce errors. The entire pipeline chart and step-by-step technical information, such as software used, time required for processing and exported file format are reported in Figure S1.

Once a single-cell suspension is prepared from the organism of interest, the cells are stained with a combination of dyes that are expected to function independently irrespective of the species used, and which have high affinity for specific cellular organelles such as nuclei, or molecules associated with metabolic states such as reactive oxygen species. We validated reagents experimentally by determining that nuclear dyes stain intracellular material matching expected characteristics of nuclear DNA or by activation of cells with drugs to change their metabolic state. The labelled samples are then run on the ImageStream®^X^ Mark II (Amnis Millipore Sigma) and images of individual events are collected for each channel of interest (Figure 1A).

Feature values from both morphological and fluorescent data, such as cell size and nuclear size, are extracted from the cell images using IDEAS software (Amnis Millipore, free for download upon creation of Amnis user account) (Figure 1B, Table S1a and S1b for feature description). Correlation between features is calculated and redundant features are trimmed as well as samples that, among replicates, are outliers (Figure S2 and S3). This prevents clustering artifacts potentially caused by having multiple features providing the same information or including samples that are not representative (Figure 1B). Finally, fluorescence intensity features are transformed to improve homogeneity of variance of distributions and, if used, DNA staining is normalized to remove intensity drift between samples and thus align the 2N and 4N DNA content histogram peaks (Figure 1B and S4).

Exported feature quantifications are used for clustering the events. Dimensionality reduction and visualization of clusters is achieved by generating force directed layout (FDL) graphs into VorteX clustering environment (Figure 1C) (free to install) (Samusik, Good, Spitzer, Davis, & Nolan, 2016). Cell images for events within each cluster can be visualized using FCS Express together with custom R scripts (Figure 1D). These visualization tools and the cluster feature averages *(i.e.,* the mean value of each feature for each cluster) (Figure 1E) allow to explore the images of selected groups of events and the features that differ between cells belonging to separate clusters. If control and treatment samples are included, a statistical analysis using negative binomial regression to compare cell counts per cluster between samples is also available in the Image3C pipeline. This high-throughput and unbiased analysis provides a comprehensive understanding of a cell population composition at higher resolution than what is possible with traditional histological methods.

Once this pipeline is run on a first set of samples *(e.g.,* homeostatic state) and the cell clusters are defined for the tissue of interest, the images and the relative clustering IDs can be used to train a CNN classifier in an unsupervised way (Figure 1F), including the ability to score frequency of “new” cell types that do not match any of the clusters identified at homeostasis. Therefore, future experiments in the same tissue used for training the CNN classifier can be analyzed directly through the CNN (Figure 1G). This significantly reduces the number of steps and time required to process data collected from following experiments with treated conditions. An even greater advantage is represented by the fact that, in the absence of CNN, every time new experimental sets are run it would be necessary to go again through the *de novo* clustering part of the pipeline (Figure 1B-1E) and the new set of clusters would need to be re-annotated to be compared with cell population composition observed in previous experiments. Manually matching clusters between different experimental sets might be a source of mistakes, mainly if the user is not familiar with the cell types present in the sample and if specific biomarkers or pre-existing knowledge about cell types and morphology are not available. The CNN splits all the cell images in the classes defined during the training step and allows to compare the abundance of cells with same morphology among different samples without the need to cross-annotate clusters (Figure 1F and 1G). The CNN inclusion in Image3C and the reproducibility of image acquisition through image-based flow cytometry allows use of the clusters defined from one experiment *(e.g.,* homeostatic state) to setup a classifier in an unsupervised and unbiased way for later use as a reference in analyzing experimental manipulations of these cell populations. We conclude from these results that Image3C can perform *de novo* high-throughput characterizations and define specific cell type behavior or population composition in both homeostatic and experimentally perturbed cell populations across multiple experiments.

### Image3C recapitulates cell composition of zebrafish whole kidney marrow (WKM) tissue

To test whether Image3C could identify homogeneous and biologically meaningful cell populations, we used the research organism *Danio rerio*. We obtained cells from adult zebrafish WKM (location of the hematopoietic tissue) in homeostasis condition, stained them and run on the ImageStream®^X^ Mark II. We analyzed intrinsic morphological and fluorescent features, such as cell and nuclear size, shape and darkfield signal (side scatter, SSC). Feature values were extracted from each cell image and processed through our pipeline (Table S1a for feature description). Clustering by the final set of normalized and non-redundant morphological and fluorescent features produced distinct cell populations (Figure 2A, 2B, 2C and S2).

**Figure 2:**
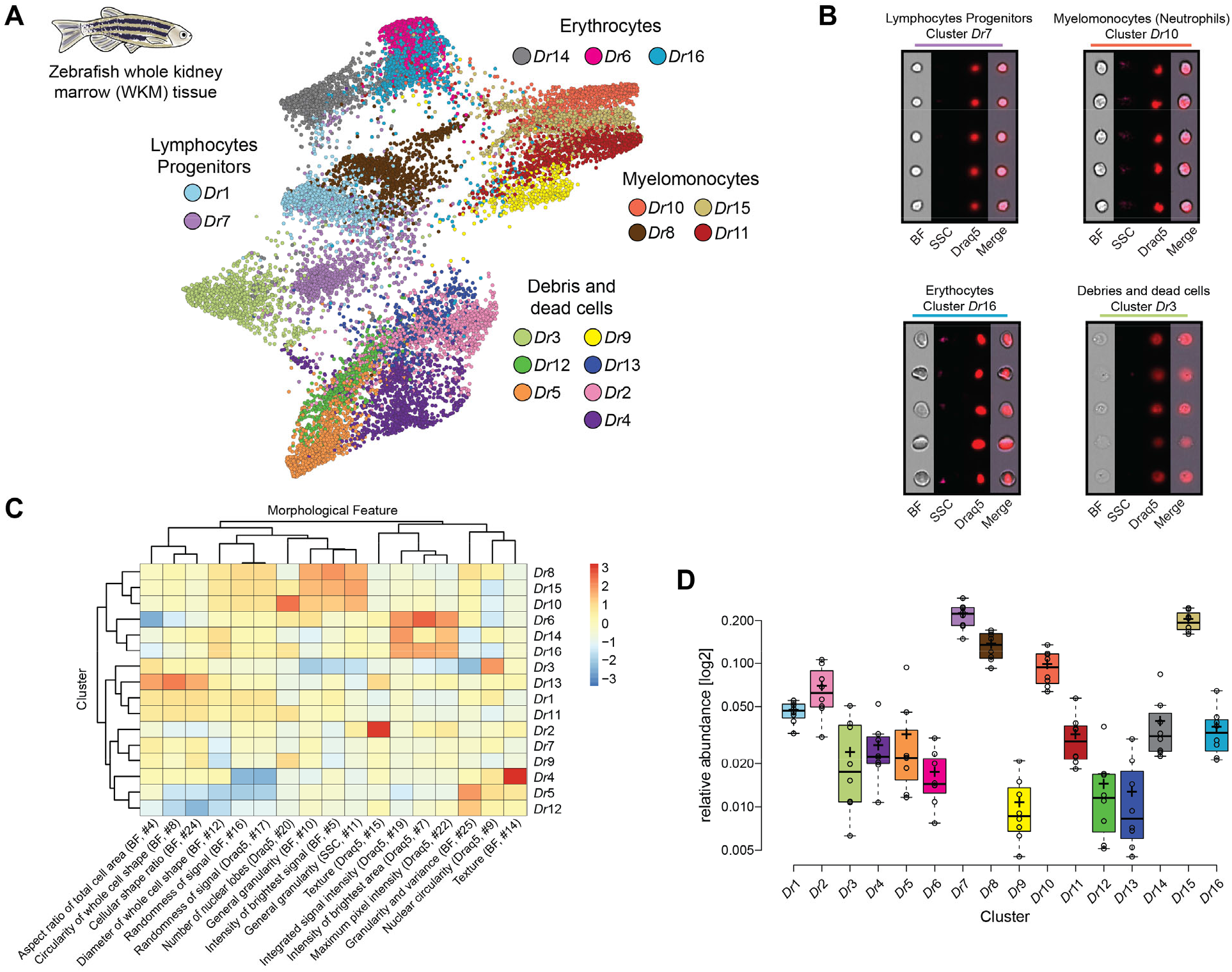
Analysis of cell composition of adult zebrafish WKM. (A) WKM tissue obtained from zebrafish is prepared for image-based flow cytometric analyses and run on the ImageStream®X Mark II (n=8). Standard gating of focused and nucleated events and manual out-gating of most erythrocytes was performed using IDEAS software (Amnis Millipore). The selected images were processed through the pipeline described in Figure 1 and clustered based only on intrinsic morphological and fluorescent feature values. FDL graph visualizes 16 clusters and each color represent a unique cell cluster. (B) Representative cell images belonging to each cluster are shown to evaluate the homogeneity of the cluster and determine morphology of the cells for cluster annotation (Data File S2). Merge represents the overlay of brightfield (BF), side scatter signal (SSC) and Draq5 (nuclear staining) signal. (C) Spearman’s correlation plot shows the average feature values of the images in each cluster to highlight morphological similarities and differences between events belonging to different clusters, such as cell size or cytoplasm granularity (Table S1a). (D) Box plot of relative abundance of events within each cluster follows the same color-code used in A.

Image3C can distinguish between the major classes of cells present in zebrafish WKM (Figure 2, Data File S1 and S2) that were described using standard sorting flow cytometry and morphological staining approaches (D. Traver et al., 2003). It is noteworthy that Image3C can clearly identify dead cells and debris (Figure 2A and 2B) allowing to optimize experimental protocols in order to minimize cell death and to run the subsequent analysis only on the intact, live cells. Image3C can identify cells with outstanding morphological features, such as neutrophils from other myelomonocytes (Figure 2B and 2C). Based on zebrafish neutrophil characteristics such as high granularity, high complexity and low circularity of the nuclei (Lugo-Villarino et al., 2010), this type of granulocytes can be easily distinguished. Other types of myelomonocytes, such as monocytes and eosinophils are here merged in the same cluster, since in zebrafish they share many morphological characteristics (Lugo-Villarino et al., 2010). Similarly, using only intrinsic morphological features for the clustering, different lymphocytes (B and T-cells) and hematopoietic stem cells cannot be separated from each other, but they can be clearly distinguished from the myelomonocytes (Figure 2A and 2B). Within the Lymphocytes/Progenitors fraction we find two clusters (*Dr*1 and *Dr*7) that mainly differ in cell diameter (Figure 2C). Whether this morphological difference has a biological implication needs to be studied in future experiments.

Image3C also enables the quantification of cell population (clusters or CNN classes) relative abundance, an important tool for comparing population composition across different treatment groups under different environmental conditions (Peuß et al., 2020). Here, we compared our results with previously published data to validate our method. Although a direct comparison with results from classical approaches (David Traver et al., 2003) is not possible since we gated out (removed analytically) mature erythrocytes before clustering (see Material and Methods), the myelomonocyte to lymphocyte ratio (M/L ratio = 1.59) is similar to the one obtained with classic histological approaches (mean M/L ratio = 1.35) (Figure 2D) (David Traver et al., 2003).

### Image3C identifies professional phagocytes in zebrafish WKM tissue

Next, we sought to determine whether Image3C could be used to characterize and quantify biological processes by identifying a tissue of interest and then comparing cellular composition dynamic, function, and physiological responses of specific cell types across a range of experimental conditions. Our goal was to detect statistically significant changes in cluster relative abundance between control and treated samples to gain a more detailed understanding of cell population dynamics and individual cell function.

As proof-of-concept, we performed a standard phagocytosis assay using WKM tissue from adult zebrafish. The single cell suspension was incubated with CellTrace Violet labeled *Staphylococcus aureus* (CTV-*S. aureus*) and with dihydrorhodamine-123 (DHR), a reactive oxygen species that becomes fluorescent if oxidized to report oxidative bursting following phagocytosis. As controls, we inhibited phagocytosis through cytoskeletal impairment with CCB incubation or through incubation with bacteria at lowered temperature by placing tubes on ice. Events collected on the ImageStream®^X^ Mark II were analyzed with Image3C and clustered in 26 distinct clusters using quantifications of morphological and fluorescent features (Table S1b for feature description), including nuclear staining, phagocytized *S. aureus* and DHR positivity (Figure 3A). Professional phagocytes are defined by their ability to take up *S. aureus* (CTV staining lies within the cell boundary) and induce a reactive oxygen species (ROS) response (bright DHR signal) (Rabinovitch, 1995). In zebrafish, professional phagocytes are mainly granulocytes and monocytic cells and can be discriminated from each other based on morphological differences, such as cell size, granularity and nuclear shape (Wittamer, Bertrand, Gutschow, & Traver, 2011). To compare samples incubated with CTV-*S. aureus* and the samples where phagocytosis is inhibited (CTV-*S. aureus* + CCB and CTV-*S. aureus* + Ice), we used the statistical analysis included in Image3C based on a negative binomial regression model (Figure 3B, 3C and S5 and Table S2 and S3). Statistical analyses reported clusters with differences in relative abundance between phagocytosis and phagocytosis-inhibited samples. Visualizing these clustered event images (Data File S3) while considering the values and intensities of morphological and fluorescent features for these clusters (Data File S1) allowed identification of 4 clusters of professional phagocytes: granulocytes within clusters *Dr*4_P, *Dr*12_P and *Dr*13_P and monocytic cells in cluster *Dr*21_P (Figure 3A, 3B). The morphology of cells in cluster *Dr*12_P is characteristic of phagocytic neutrophils (Figure 2B, 3A) that become adhesive and produce extracellular traps upon recognition of bacterial antigens (Palić, Andreasen, Ostojić, Tell, & Roth, 2007). Overall, the relative abundance of professional phagocytes is 5-10% (Figure 3C), which is in line with previous studies that estimated the number of professional phagocytes in WKM tissue of adult zebrafish using classical morphological approaches (Wittamer et al., 2011). It is also noteworthy that in line with other studies (Page et al., 2013), we did not observe a cluster of lymphocytes (*e.g.*, B-cells) that actively phagocytize CTV-*S. aureus* bacteria (Figure 2, Data File S3). Compared to the classical morphological approaches, Image3C allows to analyze thousands of events in a high-throughput and unbiased fashion, allowing the study of rare cell morphology and increasing results confidence and reproducibility. These results show that Image3C can successfully analyze biological processes since we were able to recapitulate the presence, cell type and frequency of professional phagocytes in adult zebrafish WKM organ.

**Figure 3:**
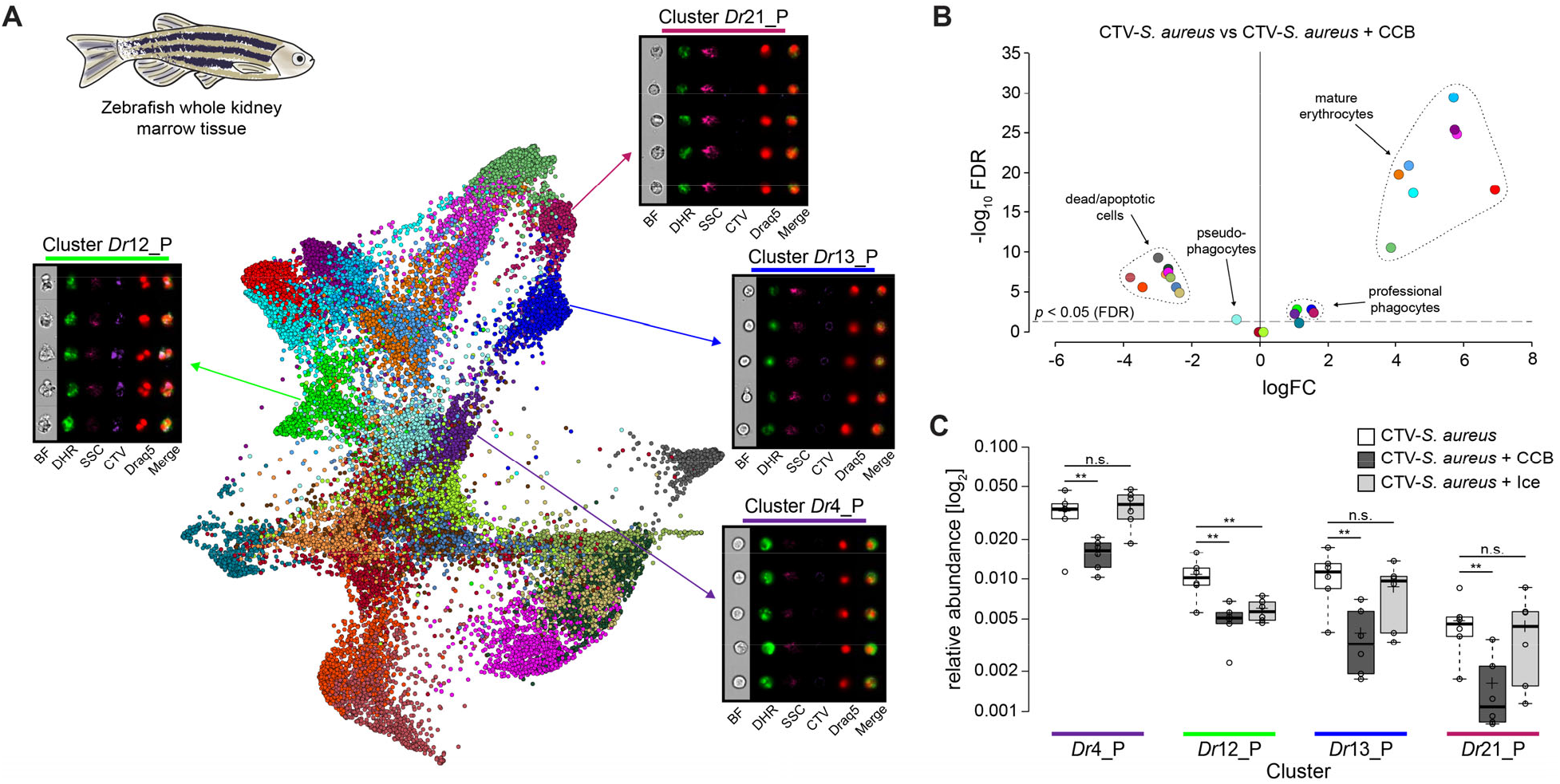
Identification of professional phagocytes in zebrafish WKM. (A) A phagocytosis assay was performed on a cell suspension obtained from zebrafish WKM tissue and the samples were subsequently run on the ImageStream®X Mark II (n=6). FDL graph shows 26 clusters and each color represents a unique cell cluster. Representative cell images belonging to the 4 clusters containing professional phagocytes are shown (Data File S3). Merge represents the overlay of DHR (ROS indicator), CTV (*S. aureus* labeling) and Draq5 (nuclear staining) channels. Table S1b reports the features used for this clustering. (B) Volcano Plot illustrates comparison of cluster relative abundance between phagocytosis samples (CTV-*S. aureus)* and inhibited-phagocytosis samples (CTV-*S. aureus* + CCB). The log fold change (logFC) is plotted in relation to the FDR (Fold Discovery Rate) corrected p-value (-log10) of each individual cluster calculated with negative binomial regression model (n=6) (Table S2). Dot color follows the same color-code used in A. (C) Box plot of relative abundances of events within the 4 clusters containing professional phagocytes. Phagocytosis samples (CTV-*S. aureus),* CCB inhibited-phagocytosis samples (CTV-*S. aureus* + CCB) and ice inhibited-phagocytosis samples (CTV-*S. aureus* + Ice) (Figure S5). Statistically significant differences are calculated using the negative binomial regression model between the phagocytosis and the inhibited-phagocytosis samples (Table S2 and S3). ** indicates p ≤ 0.01 and n.s. indicates not significantly different after FDR (n=6).

A new aspect that Image3C highlighted is that CCB selectively affects cell viability based on cell identity, introducing artifacts and cell damage, actions not specific to inhibition of phagocytosis (Figure 3B). All mature erythrocyte containing clusters had a significantly higher cell count in the CTV-*S. aureus* samples compared to the CTV-*S. aureus* + CCB ones (Figure 3B, Table S2, Data File S1). Cluster analysis revealed that erythrocytes were almost absent in samples incubated with CCB (Data File S1), while there was a significant increase in the relative abundance of clusters containing dead and apoptotic cells (Figure 3B, Table S2). Both outcomes are likely due to reduced cell viability of erythrocytes upon CCB incubation. Moreover, we excluded the possibility of higher cell death in the professional phagocytes upon CCB incubation, since pseudo-phagocytes (phagocytes with DHR response but no internalized CTV-*S. aureus)* were significantly more abundant in the CTV-*S. aureus* + CCB sample (Figure 3B, Table S2). These results are remarkable since Image3C allowed us to observe an effect of CCB on erythrocytes viability that, as far as we know, was not described before.

Image 3C analysis also uncover another important biological observation. When we inhibited phagocytosis by incubating the single cell suspension on ice (CTV-*S. aureus* + Ice) and compared the specificity of inhibition with the CTV-*S. aureus* + CCB sample (Figure 3C, Table S3), we discovered that the inhibition of phagocytosis through low temperature only affects adhesive neutrophils (cluster *Dr*_12P) (Figure 3C). This is suspected to occur as ice inhibits adhesion, while CCB effectively blocks phagocytosis in all professional phagocytes in zebrafish WKM tissue by acting on the cytoskeleton. The use of Image3C allowed us to specifically identify cell types that are sensible to low temperature and those that are not, confirming the existence of different phagocytosis mechanisms and providing additional knowledge about pro and cons of different protocols that can be applied to inhibit phagocytosis based on specific goals and needs.

### Image3C recapitulates cell composition of a freshwater snail hemolymph

Since we aim to provide a tool that is widely applicable, we tested Image3C versatility on the apple snail *P. canaliculata* an emerging organism for which molecular and cell biological tools have yet to be fully developed. As such, we repeated the same experiments done in zebrafish on the hemolymph of *P. canaliculata.* For morphological examination of the cellular composition of the hemolymph collected from adults in homeostasis conditions, we stained the single cell suspensions with Draq5 (DNA dye) and ran on the ImageStream®^X^ Mark II. We used Image3C to analyze the images of the events and we identified 9 cell clusters (Figure 4A). Two of these clusters were comprised of cell doublets, debris and dead cells (clusters *Pc5* and *Pc*8) and the other clusters, based on inspection of cell images, were grouped into 2 main categories (Figure 4A and Data File S4). The first category includes small blast-like cells (cluster *Pc*4) and intermediate cells (clusters *Pc2* and *Pc*3) with high nuclear-cytoplasmic (N/C) ratio. These cells morphologically resemble the Group I hemocytes previously described using a classical morphological approach (Accorsi, Bucci, de Eguileor, Ottaviani, & Malagoli, 2013). The second category was comprised of larger cells with lower N/C ratio and abundant membrane protrusions (clusters *Pc*1, *Pc*6, *Pc*7 and *Pc*9). Likely, these cells correspond to the previously described Group II hemocytes that include both granular and agranular cells (Accorsi et al., 2013).

**Figure 4:**
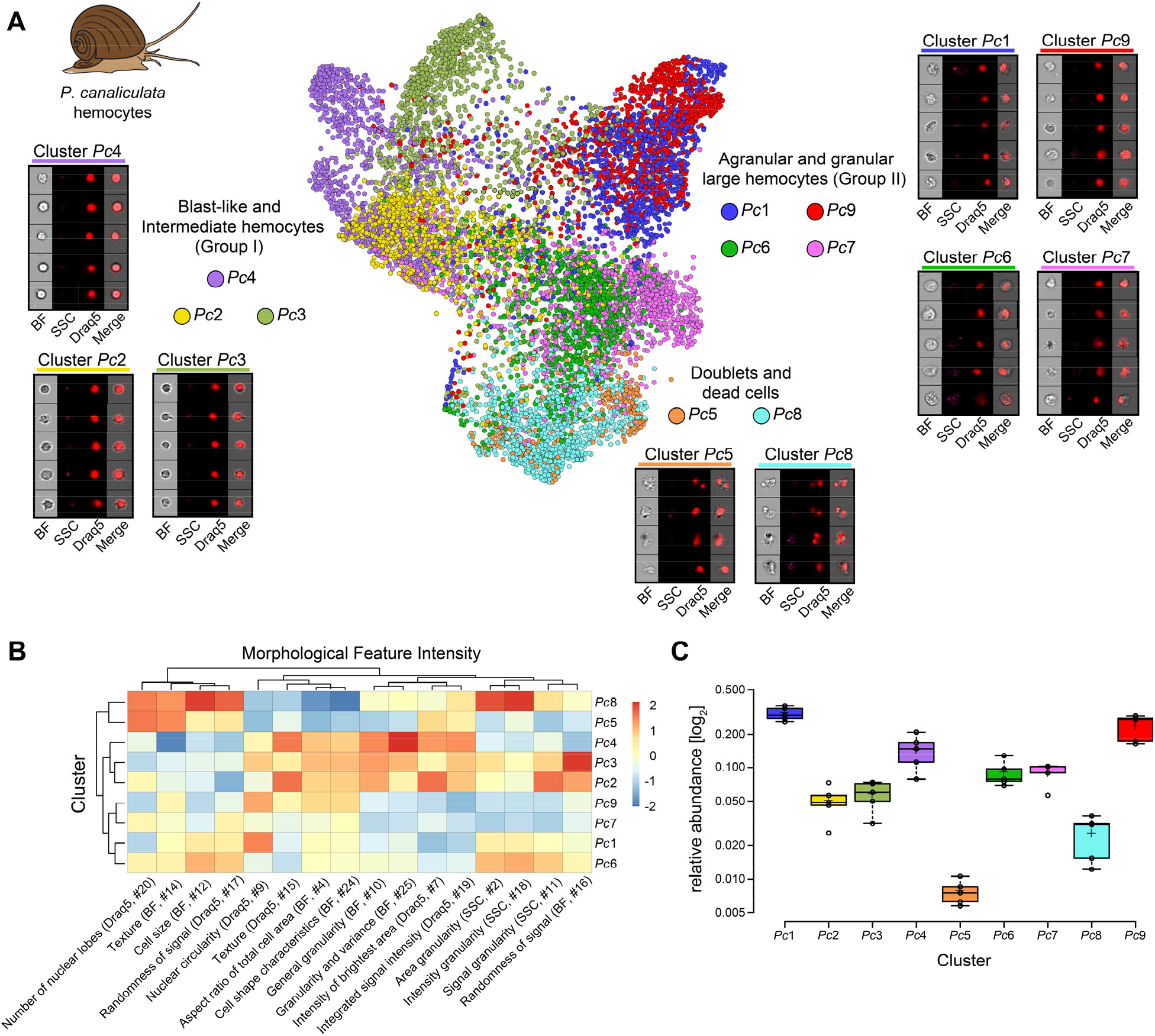
Analysis of *P. canaliculata* hemocyte population using only intrinsic morphological features of the cells. (A) Hemocytes obtained from the apple snail *P. canaliculata* are prepared for image-based flow cytometric analyses and run on the ImageStream®X Mark II (n=5). Standard gating of focused and nucleated events was performed using IDEAS software (Amnis Millipore). The selected images were processed through the pipeline described in Figure 1 and clustered based only on intrinsic morphological and fluorescent feature values. FDL graph is used to visualize the 9 identified clusters and each color represent a unique cell cluster. Representative cell images belonging to each cluster are shown to evaluate the homogeneity of the cluster and determine morphology of the cells for cluster annotation (Data File S4). Merge represents the overlay of brightfield (BF), side scatter signal (SSC) and Draq5 (nuclear staining) signal. (B) Spearman’s correlation plot shows the average feature values of the images in each cluster to highlight morphological similarities and differences between events belonging to different clusters, such as cell size or cytoplasm granularity (Table S1a). Cluster Pc6 is the one among large hemocytes with higher values in features describing cytoplasm granularity *(i.e.,* area granularity #2, intensity granularity #18 and signal granularity #11). (C) Box plot of relative abundance of events within each cluster following the same color-code used in A. Clusters *Pc*5 and *Pc*8, constituted by duplets and dead cells, are those with the lowest number of events, validating the protocol used to prepare these samples.

To identify which of these clusters were enriched for granular cells, we looked at the heatmap with feature values for each individual cluster (Figure 4B, Table S1a for feature description). Cluster *Pc*6 had the highest values for the features related to cytoplasm texture and granularity *(i.e.,* area granularity, intensity granularity and signal granularity) amongst all clusters other than cell doublets (Figure 4B, Data File S1 and S4). Based on these data, we identified cluster *Pc6* as the one containing the granular hemocytes. The clusters obtained by Image3C were not only homogeneous and biologically meaningful, but were also consistent with published *P. canaliculata* hemocyte classification obtained by classical morphological methods (Accorsi et al., 2013). Such remarkable consistency was observed both in terms of identified cell morphologies and their relative abundance in the population of circulating hemocytes (Figure 4C, Data File S4). For example, the relative abundance of the previously reported small blast-like cells is 14.0%, a value almost identical to the abundance of the corresponding cluster *Pc*4 (13.8%).

Similarly, the category of larger hemocytes, or Group II hemocytes represents 80.4% of the circulating cells as measured by traditional morphological methods, while clusters *Pc*1, *Pc*6, *Pc*7 and *Pc9* combined represent 72.4% of the events analyzed with Image3C (Figure 4C, Data File S1). A sub-set of these cells are the granular cells (cluster *Pc*6), which correspond to 7.7% of all hemocytes by classical histological methods and 8.9% by Image3C. The intermediate cells (clusters *Pc2* and *Pc*3) are less well represented in both approaches, with a relative difference in abundance of 5.6% versus 10.6% of the manually and Image3C analyzed events, respectively (Figure 4C, Data File S1). This difference is likely best explained by the remarkable difference in both the number of cells and the number of features that can be considered for analysis by Image3C. Only a few hundred hemocytes were visually analyzed using traditional histological methods based only on cell diameter and N/C ratio (Accorsi et al., 2013). In contrast, the automated pipeline used by Image3C facilitated the analysis of 10,000 nucleated events for each sample and considered 25 morphological features for each cell. The significantly higher number of morphological features simultaneously considered by Image3C also explains the higher number of clusters and improved resolution to distinguish cell types compared to the traditional methods. Hence, Image3C, not only can properly analyze cells obtained from an emerging research organism generating biologically meaningful and informative clusters but also represents an unprecedented increase in the accuracy of cell type identification over traditional histological methods, while also allowing high-throughput capability.

### Image3C identifies phagocytosis competent cells in the hemolymph of a freshwater snail

As with zebrafish, we also performed a phagocytosis experiment on hemocytes from *P. canaliculata*. Our goal was to test if it is possible with an emerging research organism to successfully discover cell phenotypes and functions and to obtain information about specific biological processes of interest by using Image3C to compare cell populations among treated and control samples.

Here, we setup the phagocytosis assay incubating the cells with CTV-*S. aureus* and DHR at room temperature. The phagocytosis was inhibited, as control, either adding EDTA (CTV-*S. aureus* + EDTA) or using low temperature by incubating samples on ice (CTV-*S. aureus* + Ice). Events collected on the ImageStream®^X^ Mark II were analyzed with Image3C and clustered in 20 distinct clusters using quantifications of morphological and fluorescent features (Table S1b for feature description), including nuclear staining, phagocytized *S. aureus* and DHR positivity (Figure 5A). We compared the phagocytosis permissive samples (CTV-*S. aureus)* with samples where phagocytosis was inhibited by EDTA incubation or low temperature using the statistical analysis included in Image3C based on a negative binomial regression model (Figure 5B, 5C and S6, Table S4 and S5). The clusters with relative abundance significantly higher in the phagocytosis samples (Figure 5B, Data File S1 and S5) and with high intensities of both DHR and bacteria signals (Figure S7), are the two clusters that we identify as enriched with professional phagocyte (cluster *Pc5*_P and *Pc17*_P) (Figure 5B and S7, Data File S5). The two clusters show a different DHR signal intensity (ROS response) from one another upon bacteria exposure (cluster *Pc5_P* with high DHR signal, cluster *Pc17_P* with low DHR signal) (Figure S7, Data File S1 and S5). Both *Pc5*_P and *Pc17*_P relative abundance is significantly higher in the phagocytosis samples compared to the EDTA treated sample (Figure 5C, Table S4), showing that EDTA inhibits phagocytosis for both types of professional phagocytes. In the sample where the phagocytosis was inhibited by low temperature, however, only cluster *Pc17*_P had a significantly lower relative abundance compared to the phagocytosis sample (Figure 5C and S6, Table S5). We can conclude that like CCB inhibition in the zebrafish phagocytosis experiment, EDTA is a more effective and generalized (not cell-type-specific) inhibitor of phagocytosis than low temperature. These results show that also in an emerging research organism, Image3C allowed discovery of new aspects of this biological process and highlighted differences among professional phagocytes that would have been difficult to detect with other methods.

**Figure 5:**
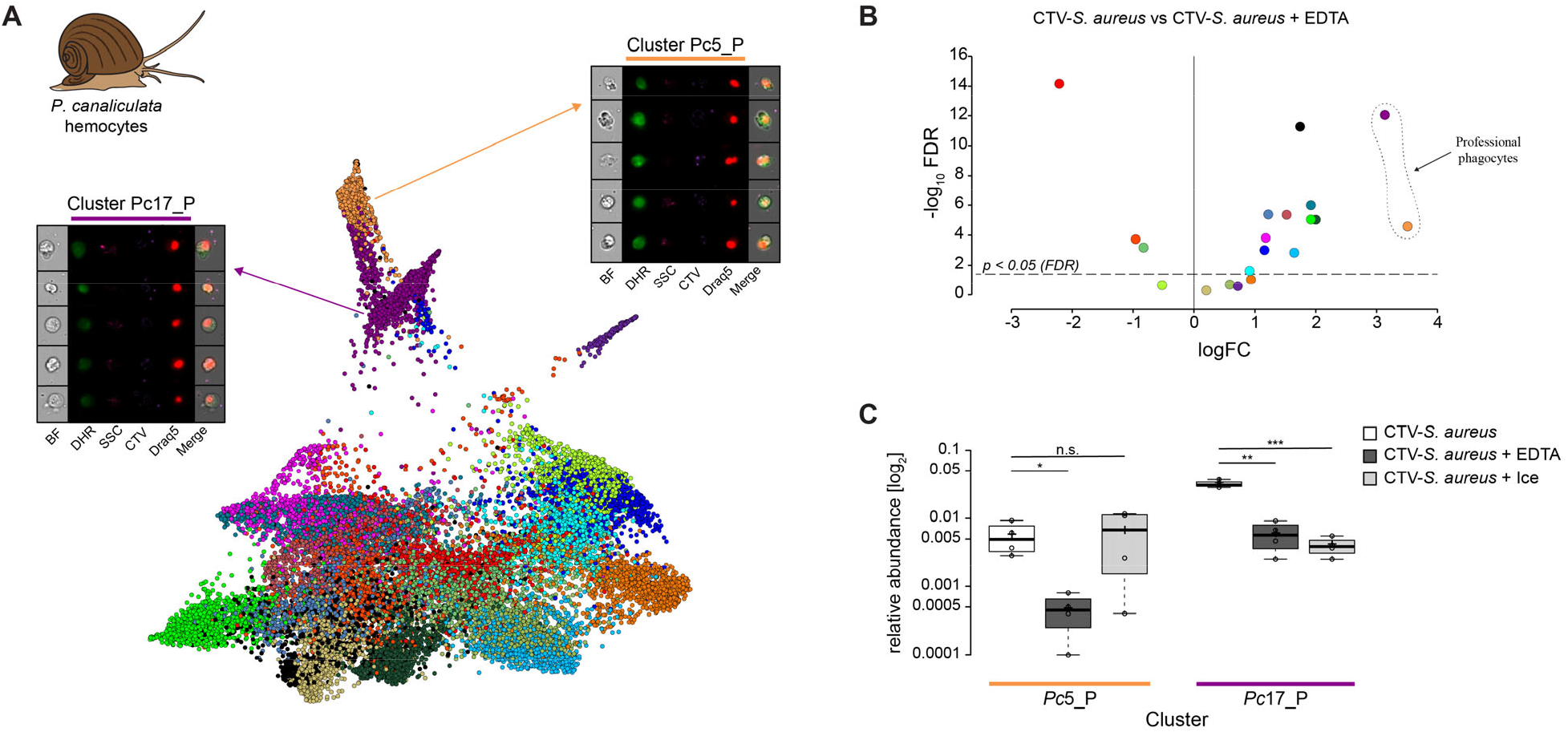
Identification of professional phagocytes among *P. canaliculata* hemocytes. (A) A phagocytosis assay was performed on a cell suspension obtained from apple snail *P. canaliculata* hemolymph and the samples were subsequently run on the ImageStream®X Mark II (n=5). FDL graph shows 20 clusters and each color represents a unique cell cluster. Representative cell images belonging to the 2 clusters containing professional phagocytes are shown (Data File S5). Merge represents the overlay of DHR (ROS indicator), CTV (*S. aureus* labeling) and Draq5 (nuclear staining) channels. Table S1b reports the features used for this clustering. (B) Volcano Plot illustrates comparison of cluster relative abundance between phagocytosis samples (CTV-*S. aureus)* and inhibited-phagocytosis samples (CTV-*S. aureus* + EDTA). The log fold change (logFC) is plotted in relation to the FDR (Fold Discovery Rate) corrected p-value (-log10) of each individual cluster calculated with negative binomial regression model (n=4) (Suppl. Table S4). Dot color follows the same color-code used in A. (C) Box plot of relative abundances of events within the 2 clusters containing professional phagocytes. Phagocytosis samples (CTV-*S. aureus),* EDTA inhibited-phagocytosis samples (CTV-*S. aureus* + EDTA) and ice inhibited-phagocytosis samples (CTV-*S. aureus* + Ice) (Suppl. Figure S6). Statistically significant differences are calculated using the negative binomial regression model between the phagocytosis and the inhibited-phagocytosis samples (Suppl. Table S4 and S5). ** indicates p ≤ 0.01 and n.s. indicates not significantly different after FDR (n=4).

The data analysis with Image3C clearly highlighted that CCB and EDTA, two classical phagocytic inhibitors commonly used in controls for phagocytosis experiments in vertebrates and invertebrates, respectively, result in a drastic change of cell morphology and cause cell death and necrosis. This consequence is not easily detectable by other methods and is therefore often overlooked. In the present work, these changes significantly modified the overall cell cluster number and distribution and indicates that the effects of CCB and EDTA on cell morphology should be taken into consideration in any study of morphological features of cells with phagocytosis properties because artifacts might be significant.

### Convolutional Neural Network (CNN) allows unbiased comparison between experiments

When determining differences between control and experimental treatments, Image3C necessarily combines images and data from all samples and then clusters the cells. This must be taken into consideration for experimental planning. Experiments meant to analyze cell composition and morphological diversity in one biological domain (*e.g.*, homeostasis condition as presented in Figure 2 and 4) should be carried out separately from those in other domains that are likely to introduce changes in the cell population composition or cell morphologies that would be a confounding factor for the *de novo* clustering in homeostasis condition. Image3C clustering works best when used, at the same time, only on samples belonging to a single experimental domain, such as homeostasis or the phagocytosis assay. An issue that emerges when analyzing different experimental sets independently is the increase of time requirement for analytical steps, the likelihood of introducing errors, and the need to repeatedly annotate the clusters in the FDL graph obtained from each experimental set. This last element is required for comparing cell type behaviors among multiple experiments and have a global understanding of their functions and response to treatments *(i.e.,* cluster #1 from one analytical run cannot be expected to match cell morphologies with cluster #1 from another run, since there is a stochastic element to the process).

This last point is probably the most challenging since mistakes can easily be introduced based on user-biases or lack of sufficient pre-existing knowledge about cell morphologies or of cell biomarkers that would allow a confident cross-annotation between multiple FDL graphs. In addition, we observed that the number of clusters drastically increases when including treatments that influence cell morphological properties of the cell. As an example, while we detected 9 unique clusters in naïve hemolymph samples, we detected 20 clusters in the phagocytosis experiment (Figure 3A and 4A). This is in part due to the fact that professional phagocytes change their morphology upon detection of pathogens (Palić et al., 2007), thus creating new clusters. Similarly, the complexity of the clustering is also increased by treatments, such as CCB and EDTA incubations, that are necessary to ensure identification of professional phagocytes, but have a strong impact on the morphology of the cells making the clustering and annotation steps more challenging and prone to mistakes since treated samples contain aberrant populations not found at homeostasis (Figure 5A, Data File S5).

To provide an alternative for streamlining the analysis of multiple experimental sets upon initial *de novo* clustering and cell type identification in homeostasis samples, we included in Image3C the possibility to use these initial images and their cluster IDs to train a CNN in an unsupervised way (Figure 1). This trained classifier can then be used to assign the cell images subsequently collected from additional experimental sets to one of the clusters defined and annotated in the homeostasis condition in a high-throughput way. In this way it will be possible to determine the behavior of a specific cell type through multiple experimental sets without re-clustering whenever new data is acquired. A crucial element that allows this approach is also represented by the ImageStream®^X^ Mark II system that provides highly reproducible and comparable images of cells coming from different experiments and acquired in different days, introducing much less variability then standard light or electron microscopy.

For our pipeline, then, a CNN (LeCun et al., 1989) based on the architecture of DenseNet (Huang, Liu, Maaten, & Weinberger, 2017) was deployed to: 1. use, as training set, images and clusters obtained from a first group of samples (*e.g.*, homeostasis conditions, naïve cells or WT samples) analyzed in an unbiased way by *de novo* clustering and 2. assign new cell images acquired through ImageStream®^X^ Mark II system to their respective classes in an unsupervised way. As proof-of-concept, we used the clusters identified for *P. canaliculata* hemocytes in homeostasis condition with the first part of the pipeline (Figure 4A) for training and setting up the CNN classifier. This approach would define the classes based on the unbiased *de novo* clustering of thousands of cells with no need for formal annotation or previous knowledge about cell types and tissue composition. To prepare a data set for training the classifier, we first combined clusters that strongly overlapped with one another in terms of morphological characteristics (*e.g.*, doublets and dead cells) to increase accuracy of the classifier (Figure 6A). We used 80% of the cells obtained in the original *P. canaliculata* dataset together with their cluster IDs to train the classifier through over 25,000 iterations. After each iteration, we tested the training with 10% of the original dataset and we determined the relative accuracy by scoring numbers of cells whose cluster ID assigned by the classifier matched the original cluster ID (Figure 6B and 6C). The remaining 10% of the original dataset was used to calculate the precision of the trained classifier. Clusters with higher support numbers obtained higher precision scores. The weighted average precision score (f1-score, precision average score across clusters controlling for support numbers) of 0.75 is relatively high considering the complexity of the phenotype (BF, darkfield and Draq5 images) (Figure 6D) and comparable to other studies using machine learning for cell classification (Blasi et al., 2016). The true probability match for each individual cell (probability for each cell that the class assigned by the classifier would match the original cluster ID) demonstrated that lower true probability matches occurred where clusters strongly overlapped or where cell phenotypes are intermediate between clusters, providing an additional layer of information about our dataset (Figure 6D).

**Figure 6:**
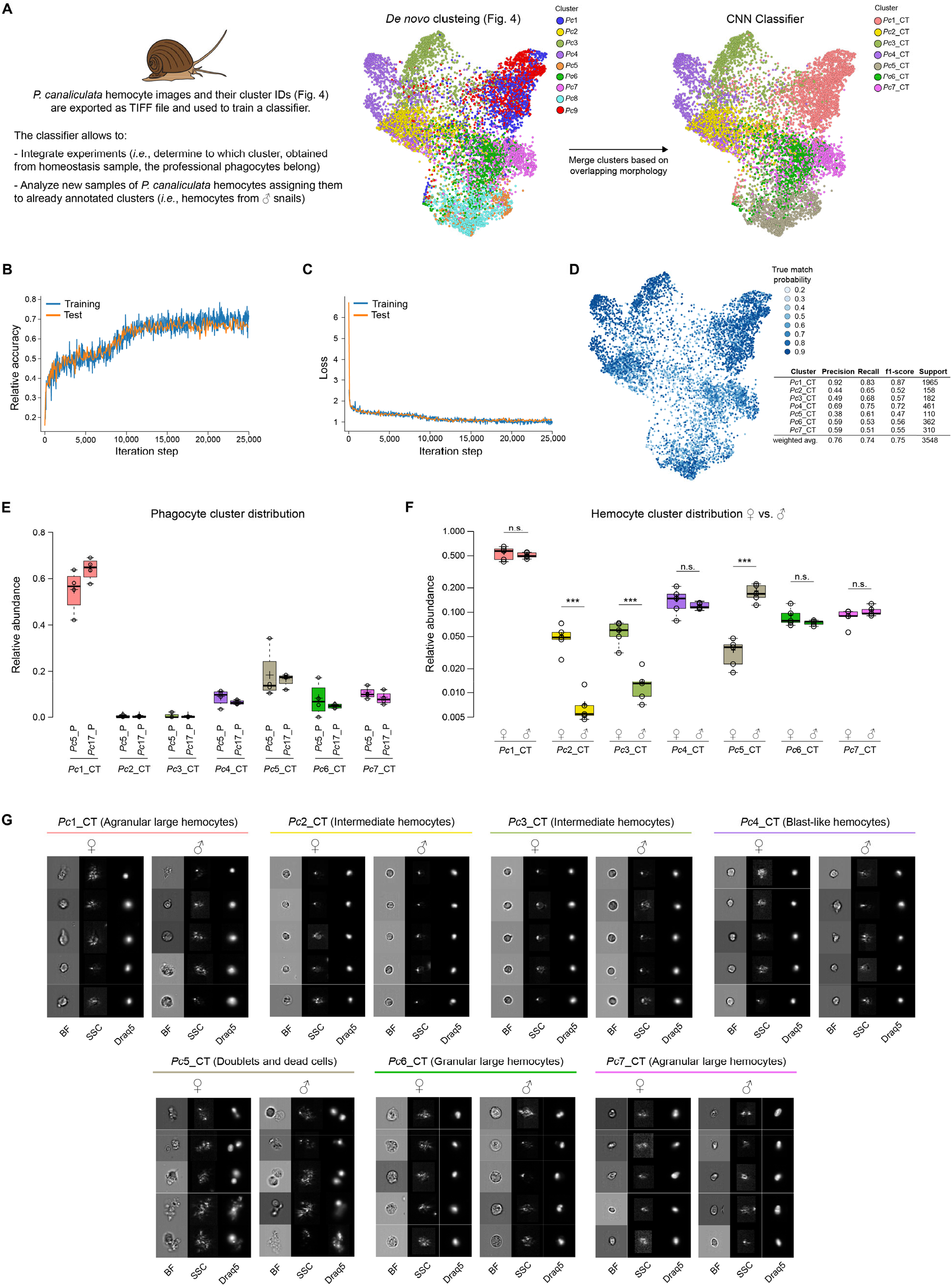
Classifier training and use of Convolutional Neural Network for integrating multiple experiments. (A) The convolutional neural network (CNN) portion of Image3C allows the unsupervised integration of multiple experiments and the analysis of additional samples assigning new events to already defined and annotated clusters. Images of cells obtained from homeostasis, naïve or WT samples and already clustered *de novo* in an unbiased way by Image3C are exported as TIFF files together with their cluster IDs. *P. canaliculata* hemocytes in homeostasis condition obtained from female apple snails (35,000 images) is the dataset used for training the CNN classifier. Before the training, clusters with cells with strongly overlapping morphology were merged (*Pc*1 was merged with *Pc*9 and named *Pc*1_CT; *Pc*5 was merged with *Pc*8 and named *Pc*5_CT). CT stands for Classifier Training. (B) The training was performed on 80% of the exported images and the testing during the training was performed on 10% of the exported images. Relative accuracy was recorded every 100 iterations for 25,000 iterations total. (C) The training was performed on 80% of the exported images and the testing during the training was performed on 10% of the exported images. Loss calculation was recorded every 100 iterations for 25,000 iterations total and indicates that the training set is not memorized. (D) The True match probability (probability that trained classifier-assigned cluster matches original cluster ID) is given for each cell of the remaining 10% of the original exported images. The detailed precision score for each cluster together with the weighted average (correcting for support) is reported in the table. (E) The CNN classifier allows for integrating experiments, such as phagocytosis assay and morphological assay, with minimal probability of introducing errors because of lack of biomarkers. The plot shows the distribution of the cells belonging to the snail professional phagocytes clusters (*Pc*5_P and *Pc*17_P from Figure 5, phagocytosis assay) among the clusters defined by morphological features using homeostasis conditions. (F) The CNN classifier allows also for the analysis of new samples obtained from the same species and the same tissue used for the training. The new events obtained running male *P. canaliculata* hemocytes, are assigned to clusters that were already annotated during the *de novo* clustering step, allowing for comparison between samples (males vs females) and statistical analysis for cluster relative abundance differences. The box plot shows the comparison of the composition of the hemocyte population between females and males (n=5). (G) Representative cell images for each cluster belonging to female and male hemocytes dataset allows to visually compare the cells used to train the classifier (female) with cells assigned by the classifier and coming from a new set of samples (males). Brightfield (BF), side scatter signal (SSC) and Draq5 (nuclear staining) signal are shown in individual channels.

To test the efficiency of this pipeline, we extracted all the images belonging to the two clusters identified in the phagocytosis assay as cluster-containing phagocytes and determined to which naïve cell-type they correspond using the CNN classifier and only the BF, SSC and Draq5 channels *(i.e.,* DHR and labeled bacteria signals where not used). We found that 59.4%, 6.2% and 9.2% of the phagocytes belonged to cluster *Pc*1_CT, *Pc*6_CT and *Pc*7_CT, respectively (Figure 6E), where CT stands for Classifier Training. These results confirmed a previously published result that used classical morphological staining and manual annotation to conclude that the hemocytes able to phagocytize were primarily Group II hemocytes (Accorsi et al., 2013). Only 8% of the phagocytes were clustered in the Group I hemocytes, here represented by cluster *Pc*2_CT, *Pc*3_CT and *Pc*4_CT, while the remaining 17.2% were assigned by the CNN to the cluster *Pc*5_CT, constituted by doublets and dead cells (Figure 6E). This result can be explained by the fact that *in vitro* phagocytosis triggers microaggregate formation (hemocyte-hemocyte adhesion) in invertebrate hemocytes that resembles the nodule formation observed *in vivo* (Walters, 1970). It is important to observe how this analysis allowed us to assign phagocytes to cell types using the annotation already performed in Figure 4A (*de novo* clustering of hemocytes in homeostasis condition) without the need to re-annotate the FDL obtained during the phagocytosis assay (Figure 5A).

To test the adaptability of the trained CNN to new datasets, we collected hemocytes from male apple snail specimens, we stained the cells with Draq5 and recorded BF, SSC and nuclei images from 10,000 cells on the ImageStream®^X^ Mark II as previously described. We extracted the images of the cells and we used our CNN classifier to determine the relative abundance of hemocytes collected from male snails in the 7 classes of the classifier (Figure 6F). First of all, we visually compared the female hemocytes clustered by the *de novo* clustering with the male hemocytes that were run on the ImageStream®^X^ Mark II at a different time and were assigned to a class by the classifier (Figure 6F). This comparison shows that the female and male hemocytes belonging to the same cluster are morphologically extremely similar and different from the hemocytes assigned to other clusters (Figure 6F). This shows that the CNN classifier can be trained with a first group of samples and then it can successfully analyze new datasets acquired later on.

The comparison between female and male hemocyte compositions revealed that the only clusters significantly different in terms of relative abundance are *Pc*2_CT and *Pc*3_CT (Group I intermediate hemocytes) and *Pc*5_CT (Figure 6F). We found *Pc*5_CT consists of dead cells and doublets and the difference in composition might be explained by variability in sample preparation and data collection. Significantly, prior studies detected no differences between females and males hemocytes composition through manual classification and counting using a classical morphological approach (Accorsi et al., 2013). The unsupervised and high-throughput analysis presented here, in contrast, allowed us to determine that both subpopulations of intermediate cells as defined by Image3C were significantly less abundant in the male animals (*Pc*2_CT: 5% and 1% in female and male, respectively; *Pc*3_CT: 6% and 1% in female and male, respectively) (Figure 6F). While the biological significance of this observation is not going to be further investigated in this paper, the discovery highlights the power of Image3C analysis compared to traditional methods for determining and quantitating the composition of cell populations. These experiments demonstrate that Image3C, in combination with the presented convolutional classifier, is capable of analyzing large experimental datasets and identifying significances with small effect sizes. Importantly, Image3C analysis is independent of observer biases and does not require prior knowledge about expected tissue composition or the expected effect of treatment on cell morphology.

## Conclusion

We have developed a powerful new method to analyze at single-cell resolution the composition of any cell population obtained from research organisms for which species-specific reagents (such as fluorescently tagged antibodies), biomarkers, single-cell atlases or a high quality genome for a scRNA-Seq approach are not available. We demonstrated that Image3C can identify cell types based on morphology and/or function through *de novo* clustering and highlight important changes in cell type abundance and cell population composition caused by experimental or natural perturbation (sex, treatment, experimental protocol). Furthermore, in combination with the CNN classifier trained on these clusters, we demonstrate that Image3C is capable of unsupervised, bias-free and high-throughput analysis of large experimental datasets making it possible to compare a specific cell type behavior or population composition across multiple experiments. Image3C is extremely versatile and can be applied to any tissue or cell population of interest and is adaptable to a variety of experimental designs. In addition, given the recent advancement in image-based flow cytometry that enables image capturing together with cell sorting (Nitta et al., 2018), a scRNA-Seq approach in combination with the Image3C pipeline would enable simultaneous analysis of both the morphological/phenotypic and genetic properties of a cell population at single cell resolution.

## Materials and Methods

### Collection of zebrafish whole kidney marrow (WKM)

Twelve-month-old, wild type, female, adult zebrafish were euthanized with cold 500 mg/L MS-222 solution for 5 min. WKM was dissected as previously described (D. Traver et al., 2003) and transferred to 40 μm cell strainer with 1 mL of L-15 media containing 10% water, 10 mM HEPES and 20 U/mL Heparin (L-90). Cells were gently forced through the cell strainer with the plunger of a 3 mL disposable syringe. The strainer was washed once with 1 mL of L-90 and the resulting single cell suspension was centrifuged at 500 rcf at 4 °C for 5 min. The supernatant was discarded, and the cells were resuspended in 1 mL of L-90 containing 5% fetal calf serum (FCS), 4 mM L-Glutamine, and 10,000 U of both Penicillin and Streptomycin (L-90 media). The cells were counted in a 1:20 dilution on the EC-800 flow cytometer (Sony) using scatter properties.

### Collection of apple snail hemocytes

Specimens of the apple snail *Pomacea canaliculata* (Mollusca, Gastropoda, Ampullariidae) were maintained and bred in captivity, in a water recirculation system filled with artificial freshwater (2.7 mM CaCl2, 0.8 mM MgSO4, 1.8 mM NaHCO3, 1:5000 Remineralize Balanced Minerals in Liquid Form [Brightwell Aquatics]). The snails were fed twice a week and kept in a 10:14 light:dark cycle. Wild type adult snails, 7-9 months old and with a shell size of 45-60 mm were starved for 5 days before the hemolymph collection (Accorsi et al., 2013). If not differently specified, female snails were used for the experiments. The withdrawal was performed applying a pressure on the operculum and dropping the hemolymph directly into an ice-cold tube. The hemolymph collected from different animals was not pooled together. The hemolymph was immediately diluted 1:4 in Bge medium + 10% fetal bovine serum (FBS) and then centrifuged at 500 rcf for 5 min. The pellet of cells was resuspended in 100 μl of Bge medium + 10% FBS. The Bge medium (also known as *Biomphalaria glabrata* embryonic cell line medium) is constituted by 22% (v/v) Schneiders’s Drosophila Medium, 4.5 g/L Lactalbumin hydrolysate, 1.3 g/L Galactose, 0.02 g/L Gentamycin in MilliQ water, pH 7.0.

### Experiment 1: Morphology Assay in homeostasis conditions

WKM cells from zebrafish were isolated as described before and plated at 4 x 10^5^ cells/well in a 96-well plate in 200 μL of L-90 media and incubated for 3 h at room temperature. Cells were stained with 5 μM Draq5 (Thermo Fisher Scientific) for 10 min and subsequently run on the ImageStream®^X^ Mark II (Amnis Millipore Sigma), where 10,000 nucleated and focused events were recorded for each sample (n=8). Erythrocytes were out-gated to enrich for immune relevant cells and to prevent over-clustering in the subsequent analysis. The latter is due to the fact that fish erythrocytes are nucleated and their biconcave shape results in different morphological feature values only depending on their orientation during image acquisition.

The *P. canaliculata* hemocytes were stained with 5 μM Draq5 (Thermo Fisher Scientific) for 10 min, moved to ice and subsequently run on the ImageStream®^X^ Mark II (Amnis Millipore Sigma), where 10,000 nucleated and focused events were imaged for each sample (n=5).

### Experiment 2: Phagocytosis Assay

*Staphylococcus aureus* (Thermo Fisher Scientific) were resuspended in PBS at the final concentration of 100 mg/ml and incubated with 5 uM CellTrace Violet (CTV; Thermo Fisher Scientific) for 20 min. Labelled bacteria were centrifuged and resuspended in PBS for 3 times to remove unbound dye and then stored at −20 °C as single-use aliquots. Cells, obtained from fish WKM or snail hemolymph and in a single cell suspension, were plated in a 96-well plate at a concentration of 4 x 10^5^ cells/well in 200 μL of medium and incubated with 2 x 10^7^ CTV-coupled *S. aureus*/well for 3 h at room temperature. As control, the phagocytosis was inhibited incubating the cells + CTV-*S. aureus* mix either on ice (for both species) or with 0.08 mg/mL cytochalasin B (CCB) for zebrafish cells or with 30 mM EDTA and 10 mM HEPES for apple snail cells (Cueto, Rodriguez, Vega, & Castro-Vazquez, 2015; Li et al., 2006). After 2 h and 30 min we added 5 μM dihydrorhodamine-123 (DHR) (Thermo Fisher Scientific) to the cell suspension to stain cells positive for reactive oxygen species (ROS) production. To control for this treatment with DHR, we incubated one aliquot of cells with 10 ng/mL phorbol 12-myristate 13-acetate (PMA) to induce ROS production. At 2 h and 50 min since the beginning of incubation with CTV-*S. aureus,* all the samples were stained with 5 μM Draq5 for 10 min. After 3 h incubation with bacteria, cells were moved and stored on ice and subsequently run on the ImageStream®^X^ Mark II (Amnis Millipore Sigma), where 10,000 nucleated and focused events were imaged for each sample (at least n=4 snail and n=6 fish) at a speed of 1,000 images/sec.

### Data collection on ImageStream®^X^ Mark II

Following cell preparation, data were acquired from each sample on the ImageStream®^X^ Mark II (Amnis Millipore Sigma) at 60x magnification, slow/sensitive flow speed (1,000 images/sec), using 633, 488 and 405 nm laser excitation. Bright field was acquired on channels 1 and 9, DHR (488 nm excitation) on channel 2, CTV-*S. aureus* (405 nm excitation) on channel 7, Draq5 (633 nm excitation) on channel 11, and SSC was acquired on channel 6. Single color controls were also acquired for each fluorescent channel to allow for fluorescence spillover correction.

### *Data analysis and* de novo *cluster identification*

A schematic representation of the pipeline with software used, format of the exported files and approximation of time required for running the individual steps is provided in Figure S1. Raw images from the ImageStream®^X^ Mark II system (RIF files, a type of modified 16 bit TIFF file) were compensated (spillover and other corrections applied), background was subtracted, and features were calculated using IDEAS 6.2 software (Amnis Millipore, free for download once an Amnis user account is created). The resulting compensated image files (CIF files) were used to quantify features for all cells and samples. Table S1a and S1b report the list of features used for each organism and for each experiment and their description. These per-object feature matrices (DAF files) were then exported from IDEAS into FCS files. Exported FCS files were processed in R (R Core Team, 2014). In order to trim redundant features that contribute noise but little new information, Spearman’s correlation values for each pair of features were calculated using all events of a representative sample and one of the features of the pair was trimmed when correlation was ≥ 0.85 for the pair (Figure S2) (Caicedo et al., 2017). The Spearman’s correlation of the mean values of remaining features per each sample were then used to identify outliers among sample replicates. Samples with correlation of mean feature values below 0.85 with the set were discarded (Figure S3). Also, while morphological features did not require any transformation or normalization, fluorescence intensity features were transformed using the estimateLogicle() and transform() functions from the R flowCore package (Ellis et al., 2018; Hahne et al., 2009) to improve homoscedasticity (homogeneity of variance) of distributions. DNA intensity features were also normalized to align all 2N and 4N peak positions and to remove intensity drift between samples (Figure S4) using the gaussNorm() function from flowStats package (Hahne, Gopalakrishnan, Khodabakhshi, Wong, & Lee, 2018). The processed data was exported from R (R Core Team, 2014) using writeflowSet() function in flowCore package (Ellis et al., 2018; Hahne et al., 2009) as CSV or FCS files, depending on downstream needs for the file output.

These processed data files were then imported into the VorteX clustering environment for X-Shift k-nearest-neighbor clustering (free to install) (Samusik et al., 2016). During the import into VorteX, all features were scaled to 1SD to equalize the contribution of features towards clustering. Clustering was performed in VorteX testing a range of k values (typically from 5 to 150), choosing a final k value using the ‘find elbow point for cluster number’ function in VorteX and confirming visually that over- or under-clustering did not occur. Force directed layout (FDL) graphs of a subset of cells obtained from each set of samples were also generated in VorteX and cell coordinates in the resultant 2D space were exported along with graphML representation of the FDL graph. Finally, tabular data (CSV files) was exported from VorteX including a master table of every cell event with its cluster assignment and original sample ID, as well as a table of the average feature values for each cluster and counts of cells per cluster and per sample.

Clustering results were further analyzed and plotted in R (R Core Team, 2014) by merging all cell events and feature values with cluster assignments and X/Y coordinates for FDL graph. Using this merged data and the graphML file exported from VorteX, new FDL graphs were created for each treatment condition using the igraph package (Csardi & Nepusz, 2006) in R (R Core Team, 2014). Statistical analysis of differences in cell counts per cluster by condition were performed using negative binomial regression of cell counts per cluster, plots of statistic results and other results were generated, and CSV files containing cell ID, sample ID, feature values, X/Y coordinates in FDL graph were exported for each sample. The subsequent use of FCS Express Plus version 6 (DeNovo software, free alternative are mentioned later in the text) allowed visualization of cell images using DAF/CIF files by cluster and by customized subsets of the FDL graphs.

DAF files were opened in FCS Express Plus software and the “R add parameters” transformation feature with a custom script was used to merge the clustering data saved in the CSV files generated above with both DAF and CIF files (feature values and image sets, respectively). This allowed to visualize image galleries of cells within each cluster and gate by features of interest on 2D plots (more traditional flow cytometry analysis) for exploring the clustering results and identifying clusters and populations of interest. FCS Express Plus is a proprietary software, but similar results can be obtained with IDEAS software where a text file with cluster IDs for each image event can be imported and the cluster information can be matched to the event images.

The full complement of R packages used includes flowCore (Ellis et al., 2018; Hahne et al., 2009), flowStats (Hahne et al., 2018), igraph (Csardi & Nepusz, 2006), ggcyto (Jiang, 2015), ggridges (Wilke, 2018), ggplot2 (Wickham, 2016), stringr (Wickham, 2010), hmisc (Harrell & Dupont, 2019) and caret (Kuhn, 2008). The GitHub repository at https://github.com/stowersinstitute/LIBPB-1390-Image3C reports a detailed description of all these processing steps and includes sample scripts and workflow files.

### Setup and use of a Convolutional Neural Network (CNN) Classifier

Once the clusters were defined with the previously described *de novo* clustering analysis, we used a CNN (LeCun et al., 1989) based on the architecture of DenseNet (Huang et al., 2017) for image classification. Out of all the events captured with the ImageStream®^X^ Mark II system, we selected only single nucleated objects applying gates on area vs aspect ratio plot and Draq5 intensity plot to achieve this selection, respectively. For these objects, we exported 16 bit TIFF images (one channel per fluorescence/BrightField image ‘color’) using IDEAS 6.2 software (Amnis/Millipore, free for download once an Amnis user account is created).

Because images from the ImageStream®^X^ Mark II system have non-uniform sizes, each image was cropped or padded to 32×32 pixels using NumPy indexing (Walt, Colbert, & Varoquaux, 2011) in a Python script. The CNN consists of three dense blocks that transition from three-channel image input of 32×32×3 to a final size of 4×4×87 with 87 feature maps. A dense block includes three convolution layers, each followed by leaky ReLU activation. The last step of the block is a strided convolution used to down-sample the width and height of the feature maps by a factor of 2. The final layer of the CNN flattens the 4×4×87 array into a 1D vector of length 1392 and is fully connected to the output layer that is a 1D vector with a length of the number of classes for prediction. The CNN used softmax cross entropy for the loss function with L2 regularization, and the Adam optimizer (Kingma & Ba, 2015) was used to minimize the loss function. The CNN was implemented in Python using the TensorFlow platform (Abadi et al., 2015) and the SciPy ecosystem (T. Oliphant, 2006; T. E. Oliphant, 2007; Pedregosa et al., 2011).

The CNN was used to train a classifier using over 35,000 images of *P. canaliculata* hemolymph cell types in homeostasis condition acquired with the ImageStream®^X^ Mark II system. The event images were randomly split in 80% for training, 10% for testing during training, and 10% for final validation. The first 80% of the images were used together with their cluster IDs obtained by the *de novo* clustering to train the classifier. The learning rate for the Adam optimizer was set to 0.0006 with a decaying learning rate starting at 0.001 and decreasing by 1% each step. The training proceeded for 25,000 iterations with a batch size of 256 randomly selected images for each iteration. After each iteration, 10% of the cells of the original *P. canaliculata* dataset was used to test the classifier. The loss and accuracy of the CNN were recorded after every 100 iterations to monitor the performance. The CNN loss was defined by the softmax of the cross entropy (Dahal, 2017) between the final output and the one-hot-encoded image labels. To avoid the CNN memorizing the training set, L2 regularization was applied to the weights. The training and test set follow the same accuracy and loss trends over all iterations indicating the training set is not memorized and can generalize to predict the test set.

The finally trained classifier was tested on the remaining 10% of images that were completely new for the CNN. The trained model was saved for future use, so new images can be inferred by the network to predict cell types. The inference is very fast because only one forward pass is made through the network and no back propagation occurs. The result of the inference is a vector with length equal to the number of cell type classes. Each element of the vector will be the probability of the cell belonging to the corresponding class; the sum of the vector must be 1. Inferring a complete experiment will provide a probability vector for each image; the list of probability vectors can be saved as CSV file. For new images, the inference results will need to be examined to ensure the predictions are reliable. A large majority of the probability vectors should have a maximum greater than 0.5, and a subset of the images should be visually inspected to verify proper class assignment. The GitHub repository at https://github.com/stowersinstitute/LIBPB-1390-Image3C reports a detailed description of all these processing steps.

### Statistical Analysis

Negative binomial regression was performed on tables of cell counts per cluster and per sample and plots were generated using R (R Core Team, 2014) with the edgeR package (Robinson, McCarthy, & Smyth, 2010), which was developed for RNAseq analysis, but includes generally applicable and user-friendly wrappers for regression and modeling analysis and plotting of results. For the comparison of cellular hemolymph composition between females and males of *P. canaliculata* a one-way ANOVA with subsequent FDR (Benjamini-Hochberg) was used.

### Animal experiment statement

Research and animal care were approved by the Institutional Animal Care and Use Committee (IACUC) of the Stowers Institute for Medical Research.

## Acknowledgements

We acknowledge Hua Li for her assistance on the statistical analysis and we also thank the Laboratory Animal Services and the Aquatics Facility at the Stowers Institute for Medical Research for animal husbandry. We would like to thank Blair Benham-Pyle, Carolyn Brewster and Viraj Doddihal as well as Barbara Milutinović for their critical comments on an earlier version of this manuscript. This work was supported by institutional funding to ACB, CW, ASA and NR. ASA is a Howard Hughes Medical Institute Investigator. NR is further supported by the Edward Mallinckrodt Foundation, NIH Grant R01 GM127872, DP2DP2AG071466 and NSF IOS-1933428 and EDGE award 1923372. RP was supported by a grant from the Deutsche Forschungsgemeinschaft (PE 2807/1-1). AA was supported by the Emerging Models grant from the Society for Developmental Biology (SDB) and the postdoctoral fellowship from the American Association of Anatomists (AAA).

## Author Contributions

RP, ACB and AA conceived and designed the study with input from ASA and NR. RP performed *D. rerio* experiments. AA performed *P. canaliculata* experiments. ACB conceived and wrote the Image3C pipeline and associated R-scripts. CW designed and optimized the convolutional neural network. RP, ACB, AA and CW analyzed and interpreted the data. RP, ACB and AA wrote the paper. All authors read and edited the paper.

## Data availability statement

All original data underlying this manuscript can be accessed from the Stowers Original Data Repository at http://www.stowers.org/research/publications/libpb-1390. Image3C code and description is freely available at the GitHub repository https://github.com/stowersinstitute/LIBPB-1390-Image3C.

## Conflict of interest

The authors declare no conflict of interest.

## Supplemental Figures

**Figure S1:**
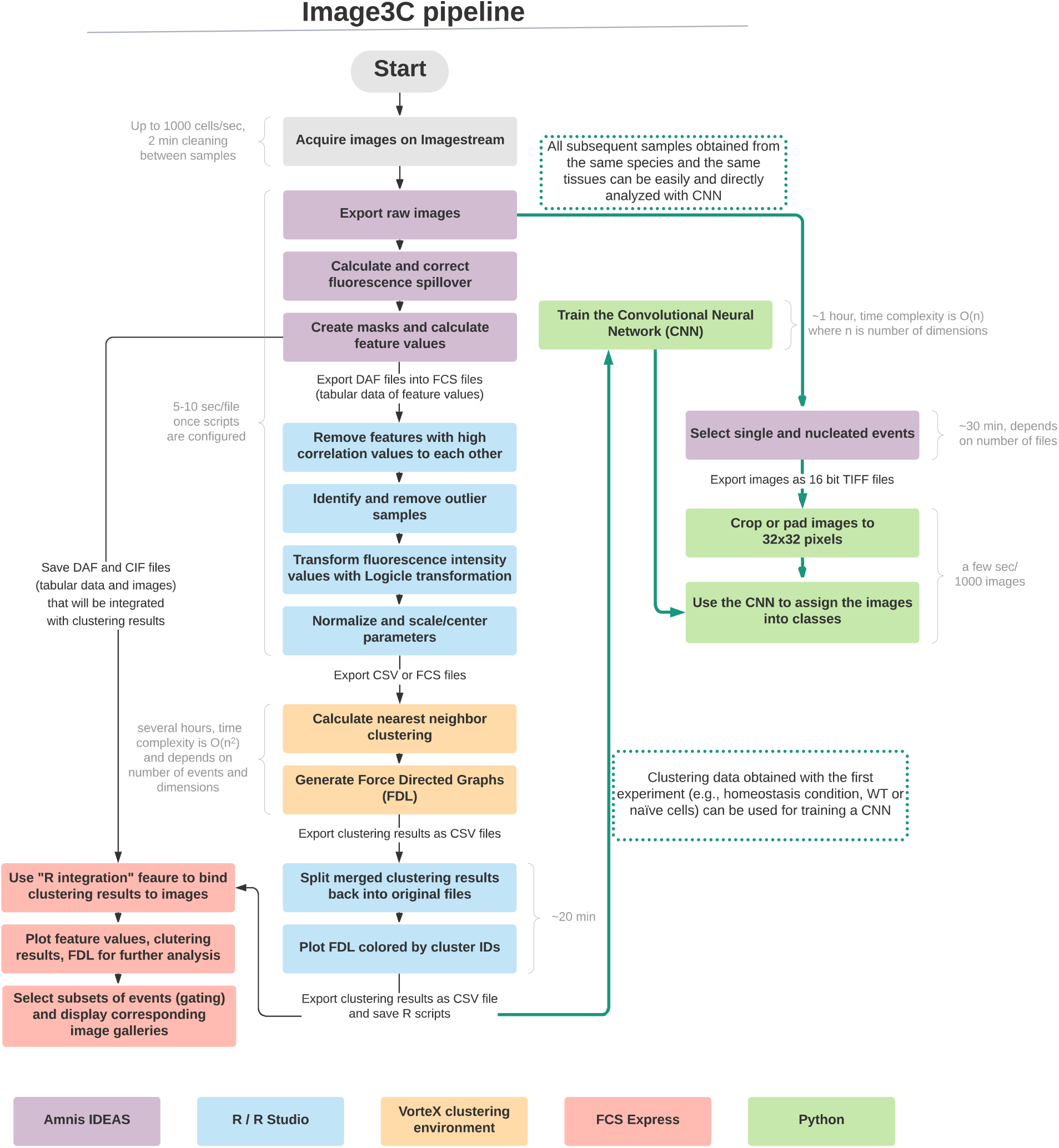
Image3C pipeline with step-by-step technical information. Once the images are collected this pipeline can be followed step-by-step. The software used are color-coded throughout the pipeline, approximation of the time needed to run the individual steps is provided in light gray. The central column and the left side represent the de novo clustering portion of the pipeline, while the right side (green arrows) represents the use of the CNN, being trained with the clusters defined in an unbiased way from the de novo clustering and able to process new samples.

**Figure S2:**
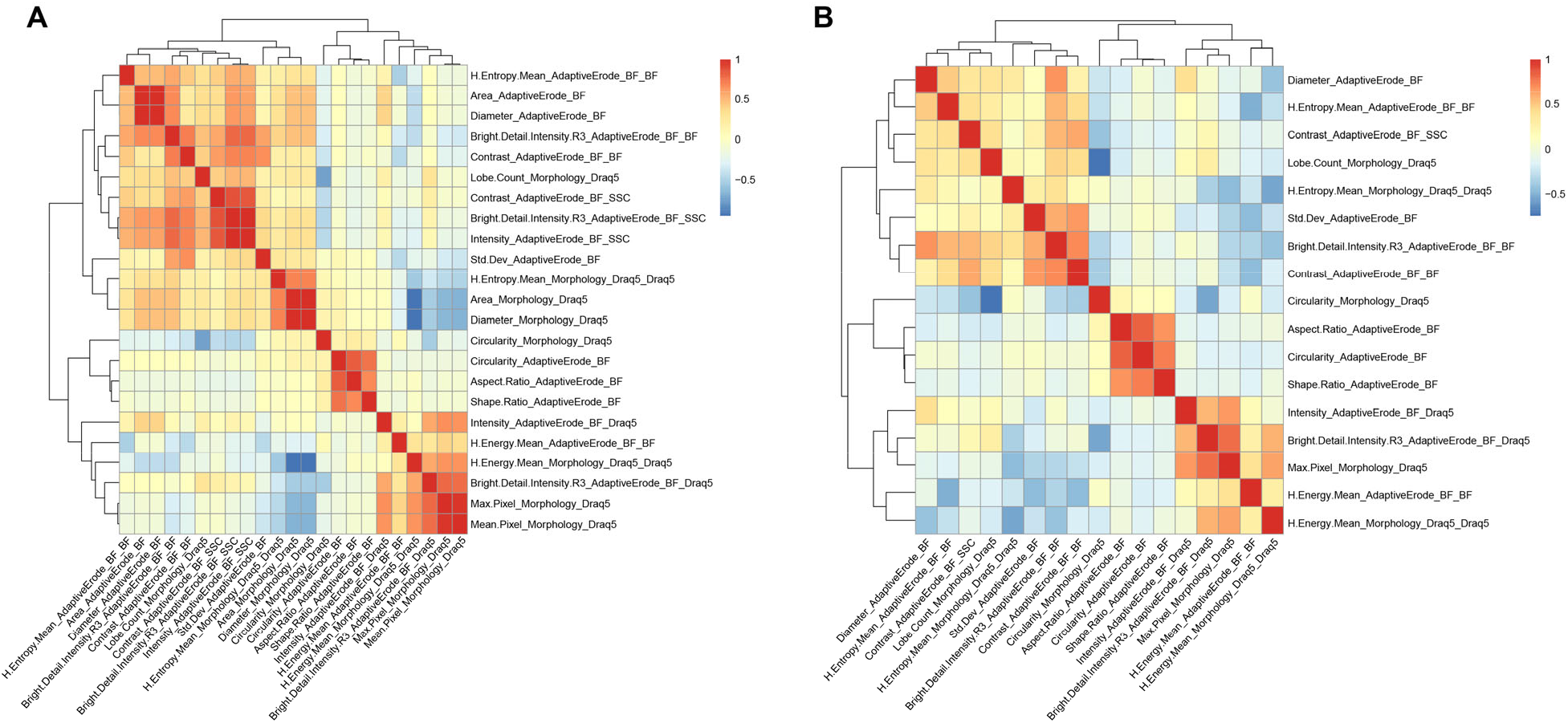
Feature correlation. An example of feature correlation and feature trimming step is shown. The correlation between feature is calculated (a) and the redundant features are removed in order to leave only one representative feature for each group of features with high correlation (b).

**Figure S3:**
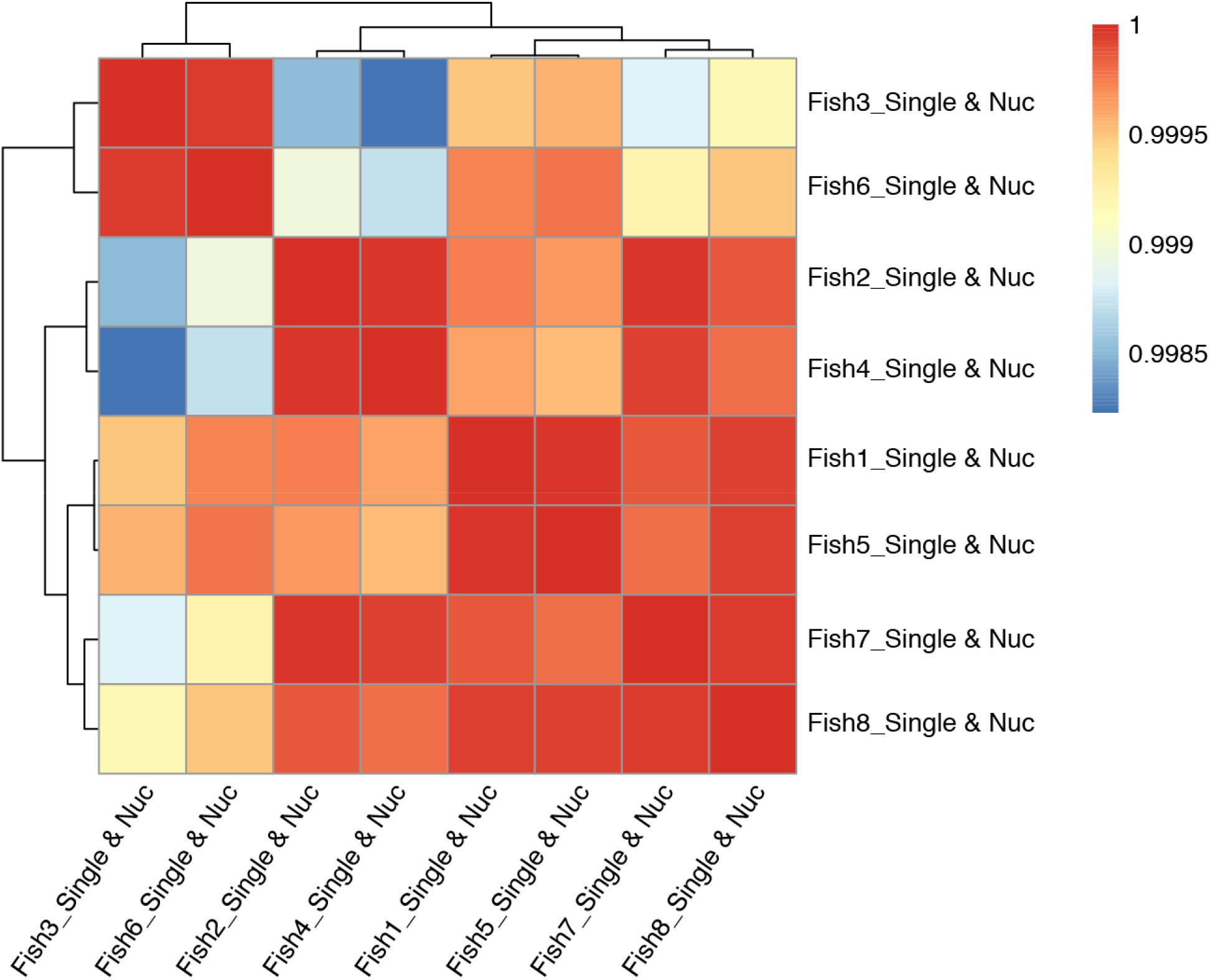
Sample correlation. An example of sample correlation and outlier sample removal step is shown. The correlation between samples is calculated and samples with clear outlier profile are removed from the subsequent analysis.

**Figure S4:**
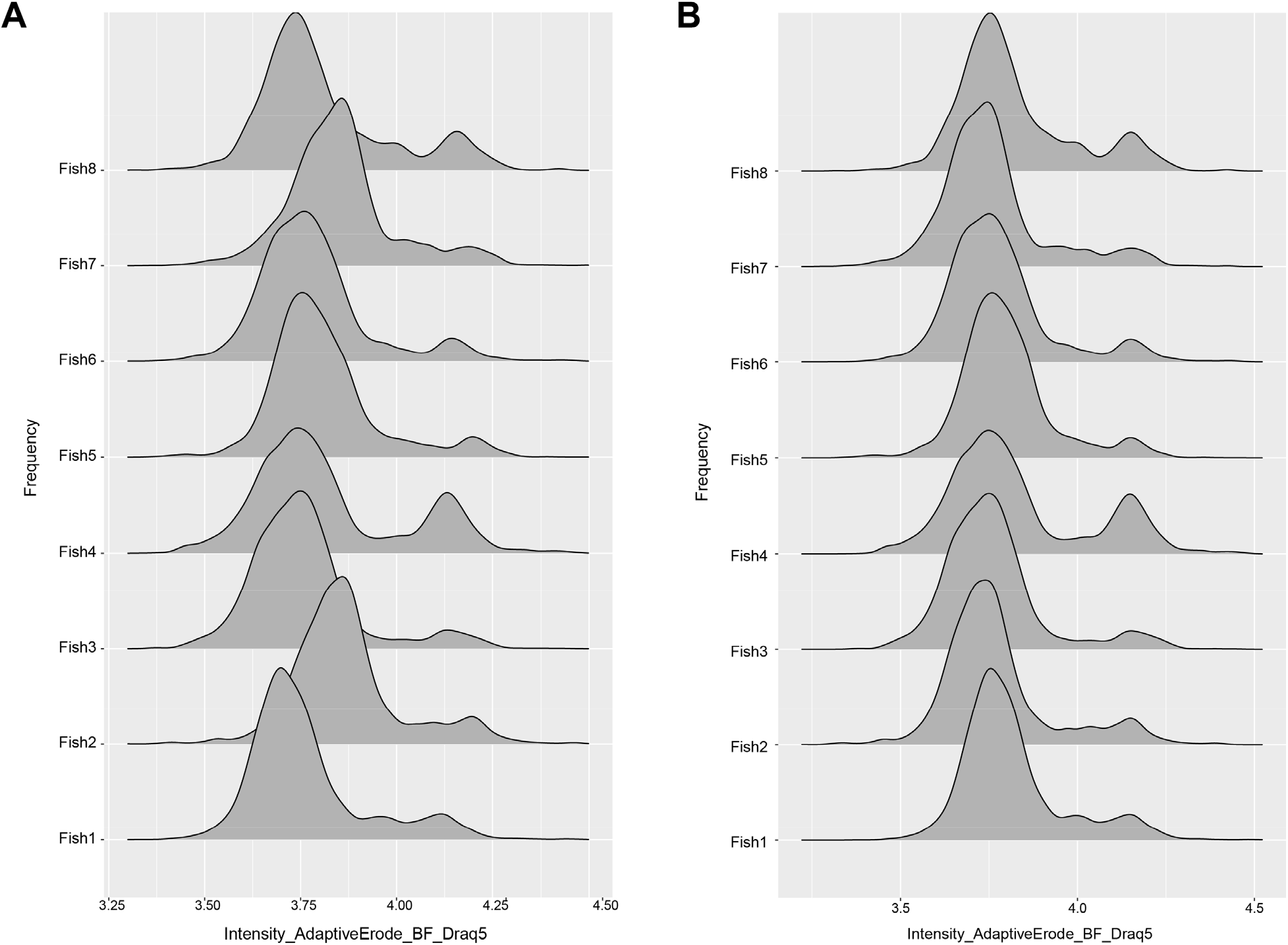
DNA normalization. An example of DNA normalization step is shown. The fluorescent peaks of DNA staining for all the samples of one experiment are shown before (a) and after (b) normalization that aims to align the 2N and 4N peaks across samples.

**Figure S5:**
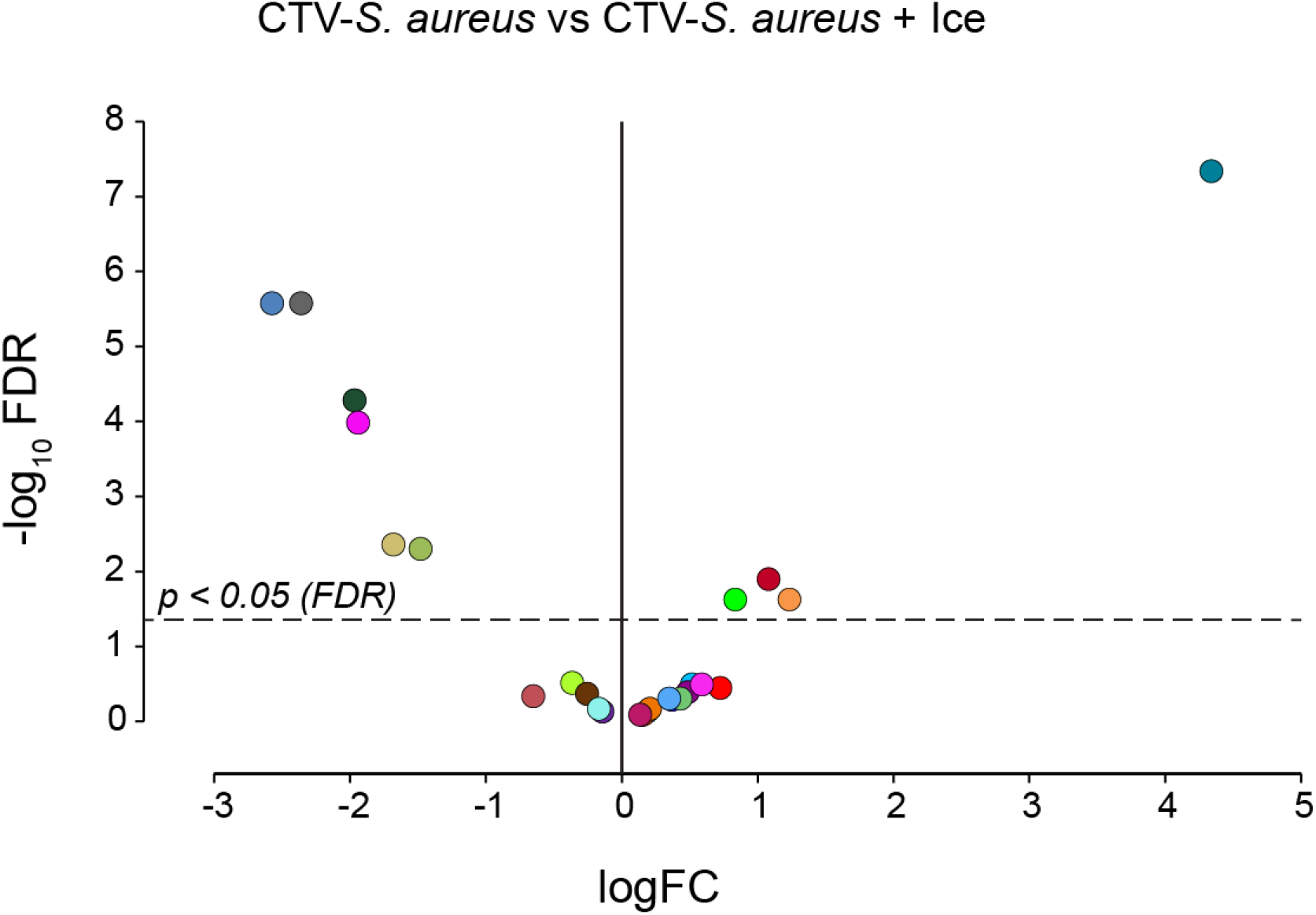
Phagocytosis assay on fish WKM with phagocytosis inhibited by ice. Volcano Plot illustrates comparison between phagocytosis samples (CTV-S. aureus) and inhibited-phagocytosis samples (CTV-S. aureus + Ice). The log fold change (logFC) is plotted in relation to the FDR (Fold Discovery Rate) corrected p-value (-log10) of each individual cluster calculated with negative binomial regression model (n=6) (Table S3). Dot color follows the same color-code used in Figure 3A.

**Figure S6:**
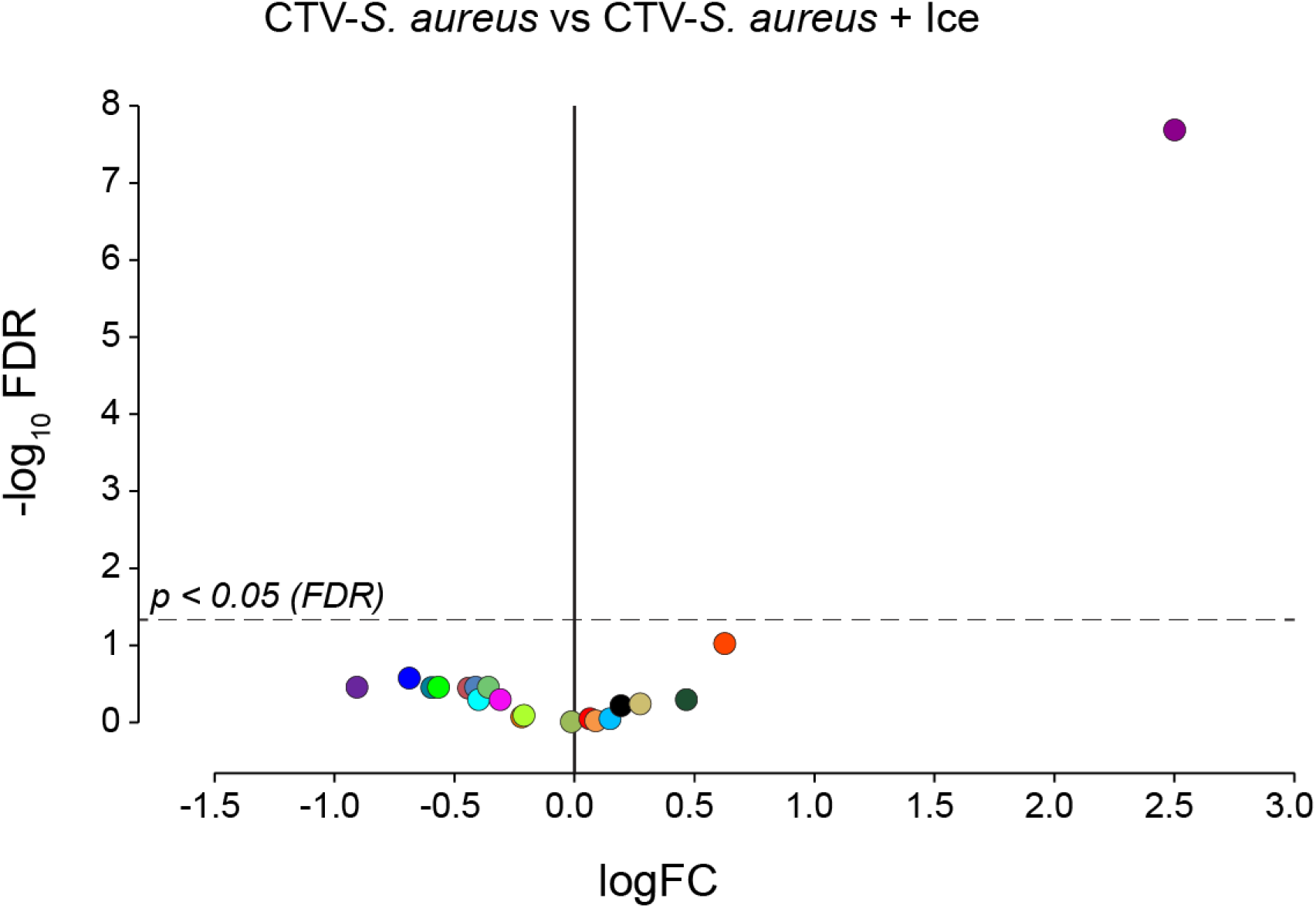
Phagocytosis assay on snail hemocytes with phagocytosis inhibited by ice. Volcano Plot illustrates comparison between phagocytosis samples (CTV-S. aureus) and inhibited-phagocytosis samples (CTV-S. aureus + Ice). The log fold change (logFC) is plotted in relation to the FDR (Fold Discovery Rate) corrected p-value (-log10) of each individual cluster calculated with negative binomial regression model (n=4) (Table S5). Dot color follows the same color-code used in Figure 5A.

**Figure S7:**
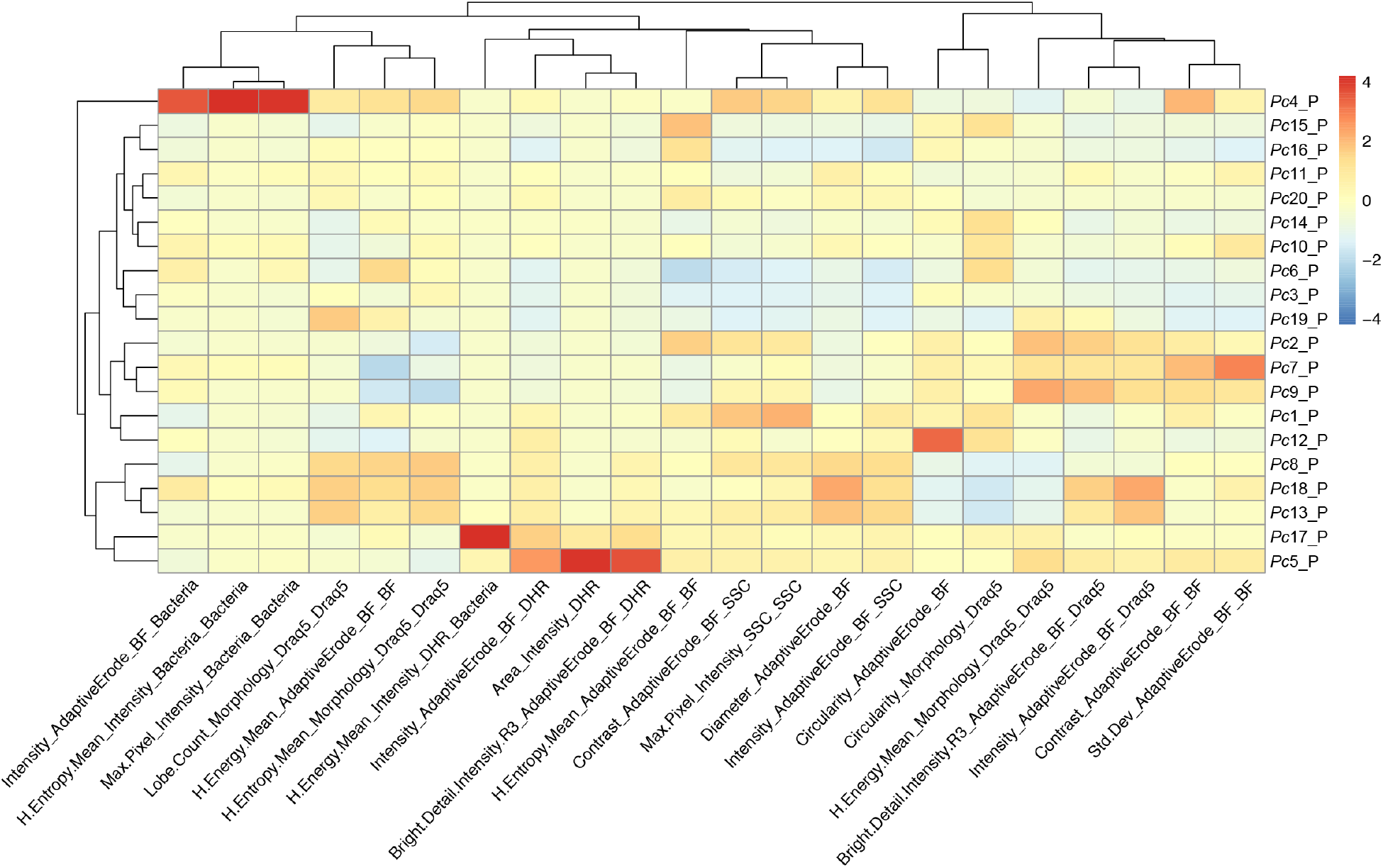
Feature heatmap after phagocytosis assay on snail hemocytes. Spearman’s correlation plot shows the average feature values of the images in each cluster to highlight morphological similarities and differences between events belonging to different clusters, such as cell size or cytoplasm granularity (Table S1b). The clusters with higher DHR signal and high bacteria are the one identified as containing professional phagocytes (*Pc*5_P and *Pc*17_P). This is also confirmed by the statistical analysis performed between the phagocytosis samples (CTV-*S. aureus*) and inhibited-phagocytosis samples (CTV-*S. aureus* + EDTA) (Figure 5B and Table S4).

## Supplemental Data Files

**Data File S1: Cell cluster properties**

Cell cluster properties (*e.g.*, cell numbers per cluster, cell feature used for clustering, feature values for each cluster) for zebrafish WKM morphology and phagocytosis assay and for snail hemocyte morphology and phagocytosis assay.

**Data File S2: Cell Gallery for zebrafish WKM in homeostasis condition**

Representative cell images belonging to each individual cluster identified by Image3C for zebrafish WKM in homeostasis condition are shown. BF is Bright Field, SSC is Side Scatter Signal and Draq5 is nuclear staining. Merge represents the overlay of BF, SSC and Draq5.

**Data File S3: Cell Gallery for zebrafish WKM after phagocytosis assay**

Representative cell images belonging to each individual cluster identified by Image3C for zebrafish WKM after phagocytosis assay are shown. BF is Bright Field, DHR is a fluorescent ROS indicator, SSC is Side Scatter Signal, Bac is CTV signal (*S. aureus* labeling) and Draq5 is nuclear staining. Merge represents the overlay of DHR, Bac and Draq5.

**Data File S4: Cell Gallery for apple snail hemocytes in homeostasis condition**

Representative cell images belonging to each individual cluster identified by Image3C for snail hemocytes in homeostasis condition are shown. Ch01 is Bright Field, Ch06 is SSC (Side Scatter Signal) and Ch11 is Draq5 (nuclear staining). Merge represents the overlay of Ch01, Ch06 and Ch11.

**Data File S5: Cell Gallery for apple snail hemocytes after phagocytosis assay**

Representative cell images belonging to each individual cluster identified by Image3C for snail hemocytes after phagocytosis assay are shown. Ch01 is Bright Field, Ch02 is DHR signal (ROS indicator), Ch06 is SSC (Side Scatter Signal), Ch07 is CTV signal (*S. aureus* labeling) and Ch11 is Draq5 (nuclear staining). Merge represents the overlay of Ch02, Ch06, Ch07 and Ch11.

## Supplemental Tables

**Table S1a:**
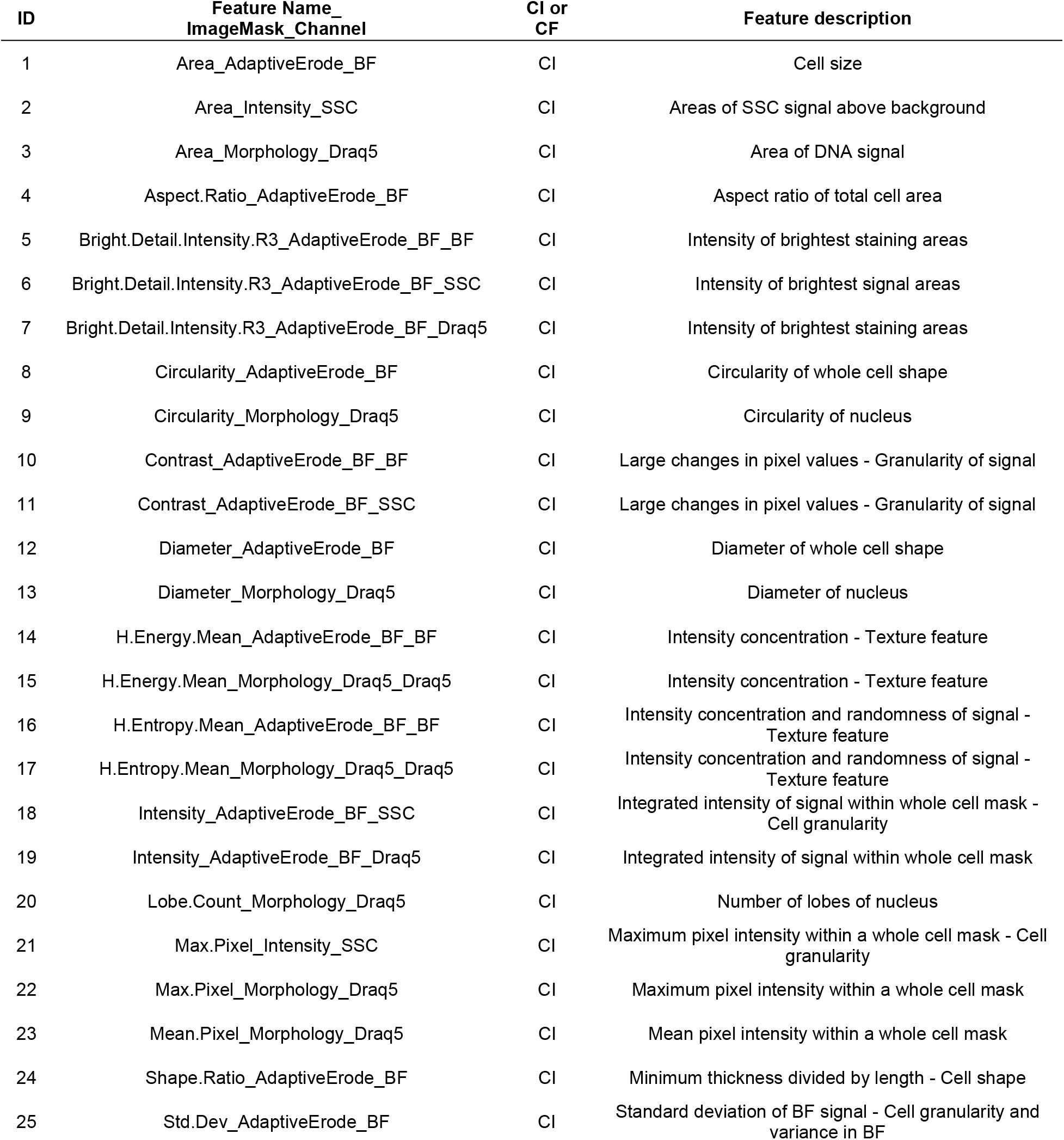
Names and descriptions of the features quantified by IDEAS software and used for clustering events based on cell morphology in the homeostasis cell composition experiment. BF is Bright Field, CI is Cell Intrinsic, CF is Cell Function.

| ID | Feature Name_<br>ImageMask_Channel | CI or<br>CF | Feature description |
| --- | --- | --- | --- |
| 1 | Area_AdaptiveErode_BF | CI | Cell size |
| 2 | Area_Intensity_SSC | CI | Areas of SSC signal above background |
| 3 | Area_Morphology_Draq5 | CI | Area of DNA signal |
| 4 | Aspect.Ratio_AdaptiveErode_BF | CI | Aspect ratio of total cell area |
| 5 | Bright.Detail.Intensity.R3_AdaptiveErode_BF_BF | CI | Intensity of brightest staining areas |
| 6 | Bright.Detail.Intensity.R3_AdaptiveErode_BF_SSC | CI | Intensity of brightest signal areas |
| 7 | Bright.Detail.Intensity.R3_AdaptiveErode_BF_Draq5 | CI | Intensity of brightest staining areas |
| 8 | Circularity_AdaptiveErode_BF | CI | Circularity of whole cell shape |
| 9 | Circularity_Morphology_Draq5 | CI | Circularity of nucleus |
| 10 | Contrast_AdaptiveErode_BF_BF | CI | Large changes in pixel values - Granularity of signal |
| 11 | Contrast_AdaptiveErode_BF_SSC | CI | Large changes in pixel values - Granularity of signal |
| 12 | Diameter_AdaptiveErode_BF | CI | Diameter of whole cell shape |
| 13 | Diameter_Morphology_Draq5 | CI | Diameter of nucleus |
| 14 | H.Energy.Mean_AdaptiveErode_BF_BF | CI | Intensity concentration - Texture feature |
| 15 | H.Energy.Mean_Morphology_Draq5_Draq5 | CI | Intensity concentration - Texture feature |
| 16 | H.Entropy.Mean_AdaptiveErode_BF_BF | CI | Intensity concentration and randomness of signal - Texture feature |
| 17 | H.Entropy.Mean_Morphology_Draq5_Draq5 | CI | Intensity concentration and randomness of signal - Texture feature |
| 18 | Intensity_AdaptiveErode_BF_SSC | CI | Integrated intensity of signal within whole cell mask - Cell granularity |
| 19 | Intensity_AdaptiveErode_BF_Draq5 | CI | Integrated intensity of signal within whole cell mask |
| 20 | Lobe.Count_Morphology_Draq5 | CI | Number of lobes of nucleus |
| 21 | Max.Pixel_Intensity_SSC | CI | Maximum pixel intensity within a whole cell mask - Cell granularity |
| 22 | Max.Pixel_Morphology_Draq5 | CI | Maximum pixel intensity within a whole cell mask |
| 23 | Mean.Pixel_Morphology_Draq5 | CI | Mean pixel intensity within a whole cell mask |
| 24 | Shape.Ratio_AdaptiveErode_BF | CI | Minimum thickness divided by length - Cell shape |
| 25 | Std.Dev_AdaptiveErode_BF | CI | Standard deviation of BF signal - Cell granularity and variance in BF |

**Table S1b:**
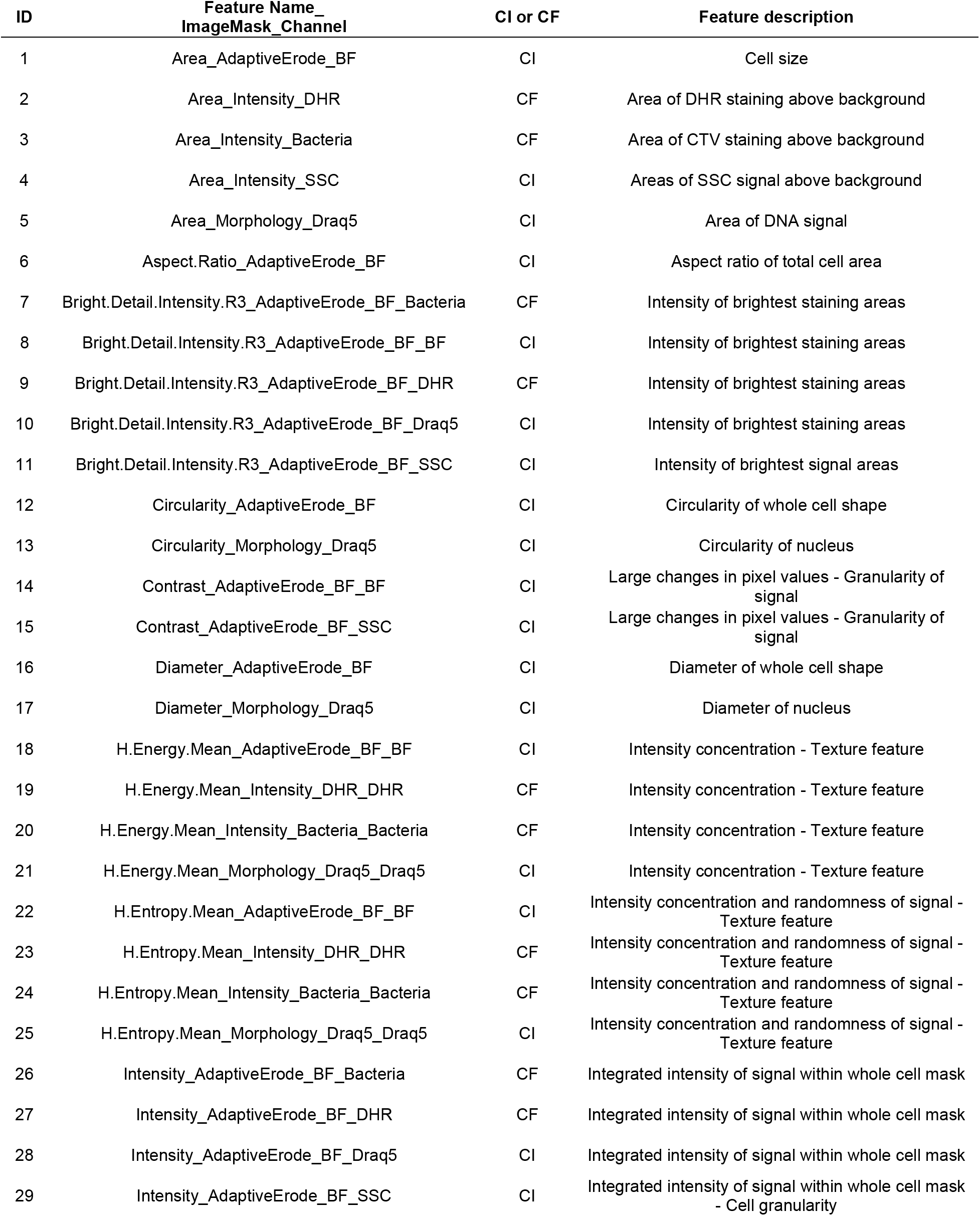
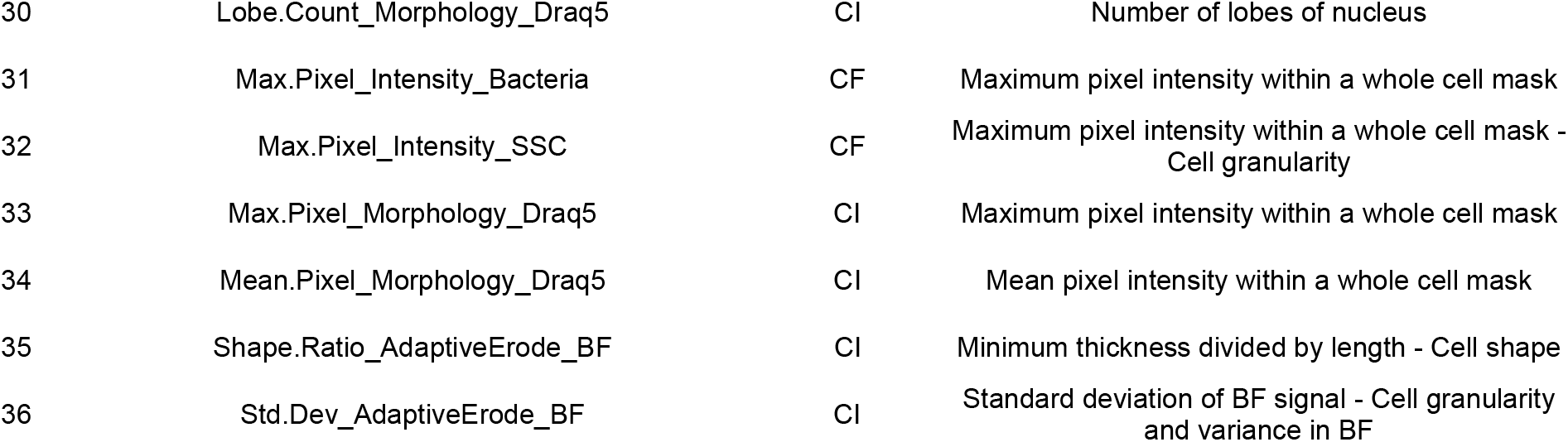
Names and descriptions of the features quantified by IDEAS software and used for clustering events based on cell morphology and function in the phagocytosis experiment. BF is Bright Field, CI is Cell Intrinsic, CF is Cell Function.

**Table S2:** Results of negative binomial regression analysis comparing cluster relative abundance between phagocytosis samples (CTV-*S. aureus*) vs phagocytosis inhibited with CCB samples (CTV-*S. aureus* + CCB) in the zebrafish phagocytosis experiment. FC is Fold Change, CPM is Count Per Million, LR is Likelihood Ratio, FDR is Fold Discovery Rate. Relative graph is reported in Figure 3B.

| Cluster ID | logFC | logCPM | LR | PValue | FDR |
| --- | --- | --- | --- | --- | --- |
| *Dr1_P* | -2.48673 | 14.76127 | 24.65067 | 6.9E-07 | 1.2E-06 |
| *Dr2_P* | -3.82090 | 15.11433 | 30.32912 | 3.7E-08 | 7.0E-08 |
| *Dr3_P* | -2.63248 | 15.10065 | 30.25504 | 3.8E-08 | 7.0E-08 |
| *Dr5_P* | -2.76060 | 14.21908 | 33.08750 | 8.8E-09 | 1.9E-08 |
| *Dr6_P* | -2.70177 | 13.37119 | 36.16033 | 1.8E-09 | 4.7E-09 |
| *Dr7_P* | -2.72126 | 14.24771 | 34.08437 | 5.3E-09 | 1.3E-08 |
| *Dr8_P* | 4.48271 | 14.88169 | 82.05466 | 1.3E-19 | 4.9E-19 |
| *Dr10_P* | -3.45902 | 14.60211 | 24.35992 | 8.0E-07 | 1.3E-06 |
| *Dr11_P* | 6.90476 | 13.91280 | 84.08534 | 4.7E-20 | 2.1E-19 |
| *Dr12_P* | 1.08728 | 12.83107 | 11.29997 | 7.8E-04 | 1.1E-03 |
| *Dr13_P* | 1.51443 | 12.94985 | 11.48213 | 7.0E-04 | 1.0E-03 |
| *Dr14_P* | -2.99602 | 11.50543 | 42.88105 | 5.8E-11 | 1.7E-10 |
| *Dr15_P* | -2.38678 | 12.70026 | 21.12804 | 4.3E-06 | 6.6E-06 |
| *Dr16_P* | 5.66338 | 14.13744 | 143.18631 | 5.4E-33 | 1.4E-31 |
| *Dr17_P* | 5.71512 | 14.80998 | 122.27043 | 2.0E-28 | 2.6E-27 |
| *Dr19_P* | 4.07753 | 14.85917 | 93.86068 | 3.4E-22 | 1.8E-21 |
| *Dr20_P* | 3.84792 | 13.05544 | 49.27037 | 2.2E-12 | 7.3E-12 |
| *Dr21_P* | 1.57131 | 11.87021 | 9.42118 | 2.1E-03 | 2.8E-03 |
| *Dr23_P* | 4.37593 | 16.67274 | 99.99839 | 1.5E-23 | 9.9E-23 |
| *Dr25_P* | 5.76977 | 13.99753 | 119.32182 | 8.9E-28 | 7.7E-27 |

**Table S3:** Results of negative binomial regression analysis comparing cluster relative abundance between phagocytosis samples (CTV-*S. aureus)* vs phagocytosis inhibited with ice samples (CTV-*S. aureus* + Ice) in the zebrafish phagocytosis experiment. FC is Fold Change, CPM is Count Per Million, LR is Likelihood Ratio, FDR is Fold Discovery Rate. Relative graph is reported in Figure S5.

| Cluster ID | logFC | logCPM | LR | PValue | FDR |
| --- | --- | --- | --- | --- | --- |
| *Dr1_P* | -2.57222 | 14.76127 | 26.21074 | 3.1E-07 | 2.7E-06 |
| *Dr3_P* | -1.47929 | 15.10065 | 10.27905 | 1.3E-03 | 5.0E-03 |
| *Dr5_P* | 1.23557 | 14.21908 | 6.95840 | 8.3E-03 | 2.4E-02 |
| *Dr6_P* | -1.96681 | 13.37119 | 19.93869 | 8.0E-06 | 5.2E-05 |
| *Dr7_P* | -1.93868 | 14.24771 | 18.19453 | 2.0E-05 | 1.0E-04 |
| *Dr9_P* | 4.34094 | 11.68675 | 36.21393 | 1.8E-09 | 4.6E-08 |
| *Dr12_P* | 0.83638 | 12.83107 | 6.79951 | 9.1E-03 | 2.4E-02 |
| *Dr14_P* | -2.35912 | 11.50543 | 26.47869 | 2.7E-07 | 2.7E-06 |
| *Dr15_P* | -1.67916 | 12.70026 | 10.80552 | 1.0E-03 | 4.4E-03 |
| *Dr24_P* | 1.08190 | 16.03192 | 8.34069 | 3.9E-03 | 1.3E-02 |

**Table S4:** Results of negative binomial regression analysis comparing cluster relative abundance between phagocytosis samples (CTV-*S. aureus)* vs phagocytosis inhibited with EDTA samples (CTV-*S. aureus* + EDTA) in the apple snail phagocytosis experiment. FC is Fold Change, CPM is Count Per Million, LR is Likelihood Ratio, FDR is Fold Discovery Rate. Relative graph is reported in Figure 5B.

| Cluster ID | logFC | logCPM | LR | PValue | FDR |
| --- | --- | --- | --- | --- | --- |
| *Pc1_P* | 1.21972 | 16.42389 | 23.86393 | 1.0E-06 | 4.1E-06 |
| *Pc2_P* | 1.52130 | 16.31424 | 23.42745 | 1.3E-06 | 4.3E-06 |
| *Pc5_P* | 3.50692 | 11.96025 | 19.19534 | 1.2E-05 | 2.6E-05 |
| *Pc6_P* | 2.00062 | 13.66811 | 21.45211 | 3.6E-06 | 9.1E-06 |
| *Pc7_P* | 1.17845 | 15.71918 | 15.65951 | 7.6E-05 | 1.5E-04 |
| *Pc8_P* | 0.91220 | 14.51336 | 5.60883 | 1.8E-02 | 2.4E-02 |
| *Pc9_P* | 1.91957 | 14.24789 | 21.73770 | 3.1E-06 | 8.9E-06 |
| *Pc10_P* | -0.95771 | 16.55159 | 15.15146 | 9.9E-05 | 1.8E-04 |
| *Pc11_P* | -2.21453 | 17.04466 | 66.60728 | 3.3E-16 | 6.6E-15 |
| *Pc12_P* | 1.92022 | 13.48376 | 27.01612 | 2.0E-07 | 1.0E-06 |
| *Pc13_P* | 1.15586 | 13.78276 | 11.69042 | 6.3E-04 | 9.7E-04 |
| *Pc14_P* | 1.74206 | 17.72100 | 51.24411 | 8.2E-13 | 5.4E-12 |
| *Pc16_P* | 1.64586 | 13.23961 | 10.80603 | 1.0E-03 | 1.4E-03 |
| *Pc17_P* | 3.13469 | 13.98527 | 55.58824 | 8.9E-14 | 8.9E-13 |
| *Pc20_P* | -0.82885 | 16.67206 | 12.56812 | 3.9E-04 | 6.5E-04 |

**Table S5:** Results of negative binomial regression analysis comparing cluster relative abundance between phagocytosis samples (CTV-*S. aureus)* vs phagocytosis inhibited with ice samples (CTV-*S. aureus* + Ice) in the apple snail phagocytosis experiment. FC is Fold Change, CPM is Count Per Million, LR is Likelihood Ratio, FDR is Fold Discovery Rate. Relative graph is reported in Figure S6.

| Cluster ID | logFC | logCPM | LR | PValue | FDR |
| --- | --- | --- | --- | --- | --- |
| *Pc17_P* | 2.50037 | 13.98527 | 37.29469 | 1.02E-09 | 2.03E-08 |

## Notes

### Competing Interest Statement

The authors have declared no competing interest.

### Summary of Updates

We've added a convolutional neural network that uses Image3C cluster information and Imagestream data for classification analysis of large datasets.

